# A predictive framework for stop-loss variants with C-terminal extensions

**DOI:** 10.1101/2025.09.01.673407

**Authors:** Jihoon G. Yoon, Sojin Lee, Yoonjung Kim, Heon Yung Gee, Min Goo Lee, Kyung-A Lee

**Affiliations:** Department of Laboratory Medicine, Gangnam Severance Hospital and Yonsei University College of Medicine, Seoul, Republic of Korea; Department of Pharmacology, Yonsei University College of Medicine, Seoul, Republic of Korea

**Author notes:** To whom correspondence should be addressed. Correspondence may also be addressed to Kyung-A Lee.

## Abstract

Stop codons dictate translation termination, and variants occurring at these sites can result in stop-loss variants, leading to C-terminal extensions with potentially significant functional consequences. Despite their clinical relevance, existing prediction tools—primarily developed for missense variants—lack sufficient accuracy for assessing stop-loss variants, mainly due to their insufficiency in accounting for the sequence features of the extended peptide. To address this gap, we developed TAILVAR (Terminal codon Analysis and Improved prediction of Lengthened VARiants), a machine-learning classifier that integrates multi-omics features spanning transcript- and protein-level properties, along with variant effect annotations. Our analyses showed that transcripts lacking downstream stop codons in the 3’ untranslated region exhibit lower evolutionary constraints. Additionally, we observed that deleterious variants exhibit greater C-terminal hydrophobicity, which is associated with reduced protein stability and increased degradation, as well as a higher aggregation propensity. TAILVAR outperformed existing benchmarks, demonstrating the highest correlation with functional experiments and establishing thresholds to classify variants as benign or pathogenic. This work offers a systematic framework for interpreting stop-loss variants, providing precise predictions of elongated protein effects that may aid genetic diagnosis and facilitate the discovery of novel disease-associated genes.

**Graphical abstract:** 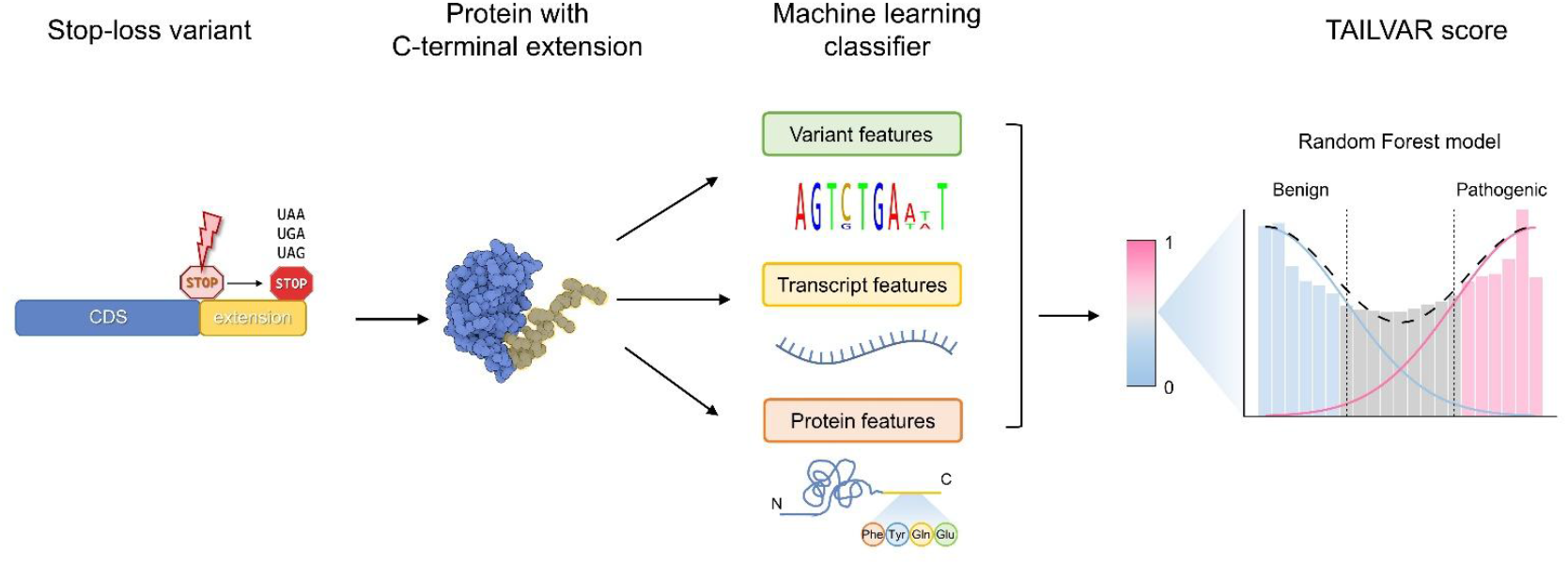

## Introduction

Protein synthesis, a fundamental biological process, is intricately governed by the genetic code, where triplet codons—comprising 64 possible combinations of the four nucleotides—play a crucial role. These codons dictate the initiation, elongation, and termination phases of messenger RNA (mRNA) translation. This translation process is tightly regulated by various factors, including eukaryotic initiation factors, eukaryotic elongation factors, and eukaryotic release factors (eRFs), which function within the 60S and 40S subunits of the ribosome (1). Specifically, eRFs recognize the three stop codons (TAA, TAG, and TGA) at the A site of the ribosome, facilitating the release of the nascent polypeptide chain from the peptidyl-transfer RNA located at the P site within the peptidyl-transferase center (2).

During the translational process, read-through events at canonical stop codons can occur under specific physiological conditions, such as oxidative stress, even in otherwise healthy tissues (3, 4). A recent study demonstrated that proteins subject to read-through are often highly expressed and enriched for intrinsically disordered C-terminal domains (5). Such events are thought to be under selective pressure across genomes, thereby mitigating potential toxic effects of non-canonical protein expression (6, 7). Beyond these physiological contexts, genetic variants at stop codons can similarly induce read-through events. These variants can be broadly categorized as stop-retained or stop-loss variants (8). While the terminal codon is changed to other stop codons, stop-retained variants generally maintain the protein’s original structure and function, resulting in minimal biological impact. Conversely, stop-loss variants replace the original stop codon with a coding amino acid, potentially leading to the translation of an extended sequence until a new stop codon is encountered in the 3’ untranslated region (UTR), if present. In cases where no stop codons are present downstream, nonstop decay (NSD), a cellular surveillance mechanism, may degrade the affected mRNA (9).

During variant interpretation, predicting the functional consequences of stop-loss variants remains challenging due to the lack of a robust framework. According to the American College of Medical Genetics and Genomics (ACMG) and the Association for Molecular Pathology (AMP) criteria (10), stop-loss variants are categorized under the PM4 criteria, representing moderate evidence of pathogenicity. A recent study indicates that the length of the C-terminal extension is a crucial determinant of pathogenic potential, with extensions exceeding 30 amino acids predicting higher risks (11). However, relying solely on extension length to predict functional outcomes demonstrated inadequate sensitivity and specificity. Furthermore, despite numerous in silico tools accurately assessing missense and other variant types, their performance in evaluating stop-loss variants remains suboptimal (12–22). This gap emphasizes the urgent need for specialized frameworks and tools to enhance the interpretation and understanding of the functional impacts of stop-loss variants.

Recently, integrating proteomic context with deep learning technologies has significantly improved the pathogenicity prediction of missense variants (23, 24). Similarly, we considered transcript and protein features of C-terminal extensions and developed a machine-learning classifier, TAILVAR (Terminal codon Analysis and Improved prediction of Lengthened VARiants), specifically to assess the pathogenicity of stop-loss variants. Our results demonstrate that TAILVAR outperforms existing tools and can be effectively applied to interpret stop-loss variants. Furthermore, to facilitate their assessment, we constructed a comprehensive database of all possible C-terminal extensions arising from single-nucleotide substitutions and indels at stop codon sites, and we lastly proposed an interpretive framework informed by our analyses.

## Materials and Methods

### Data acquisition and processing

The human reference genome GRCh38 (hg38) was utilized for analysis, which sourced from the Genomics Public Data on Google Cloud (https://console.cloud.google.com/storage/browser/genomics-public-data/resources/broad/hg38/v0/). Genomic position and nucleotide sequence information for stop codons were retrieved from GENCODE (Release 48; GRCh38.p14) (25). Coding and 3’ UTR sequences were obtained from Ensembl BioMart using the ‘biomaRt’ R package (26). We focused on the canonical transcripts from the Matched Annotation from NCBI and EMBL-EBI (MANE) project that terminated with one of the three standard stop codons (TAA, TAG, and TGA) (27). Variants were annotated using the Variant Effect Predictor (VEP) v114 (28). Stop-loss variants were filtered based on the VEP annotations, specifically targeting those categorized as ‘stop_lost’ in the “Consequence” column. Computational prediction and conservation scores were retrieved from the dbNSFP database (v5.2a) (29). The length and composition of the C-terminal extensions were determined by identifying the nearest in-frame stop codon within the 3’ UTR, translating the extended sequences, and calculating the distance from the original stop codon. Transcripts with no available 3’ UTR sequences were excluded from the analyses. The probability of being loss-of-function (LOF) intolerant (pLI) score and LOF observed/expected upper bound fraction (LOEUF) were used as gene-level constraint metrics, and they were obtained from Genome Aggregation Database (gnomAD v4.1.0) (30). To investigate the unique features of transcripts, we also analyzed mRNA half-lives. Due to the variability in mRNA half-life estimates across different measurement methods and cell types, we leveraged the Z-scores from the Saluki dataset, which integrates 39 human transcriptome-wide mRNA decay rate datasets (31). Additionally, gene essentiality classifications were sourced from DepMap.org, following the same method described previously with a more recent database (23). Specifically, gene essentiality annotations were retrieved from the DepMap Public 24Q4 dataset. Furthermore, the list of manually curated intrinsically disordered proteins (IDPs) was obtained from the DisProt database (Release 2024_12) (32).

### Algorithm and feature selection

To optimize the model performance, we explored multiple machine learning algorithms, including linear regression, decision tree, support vector machine, random forest, and neural networks, using the ‘*caret*’ R package (33). Preliminary analyses revealed that the random forest model demonstrated superior performance among the tested algorithms; thus, the TAIILVAR model was built based on a random forest algorithm implemented with the ‘*randomForest*’ R package (34). For feature selection, we integrated 4 variant-level prediction and conservation scores available for stop-loss variants. These included CADD, GERP, PhyloP100, and PhastCons100 (*V*_1_ − *V*_4_) (12–15). In addition, we incorporated transcript- and protein-related features likely to influence gene expression and protein stabilization. The transcript-level features (*T*_1_ − *T*_6_) comprised 3′ UTR length, 3′ UTR GC (%), mRNA stability, and constraint metrics such as pLI, LOEUF, and *s*_het_ (30, 31, 35). Protein-related features included the length of the original protein, the length of the introduced novel C-terminal peptide, counts of each amino acid in the extended sequence, average hydrophobicity within these peptide sequences, aggregation properties, and whether the protein is curated in the IDP database (*P*_1_ − *P*_27_). Because increased hydrophobicity in C-terminal extensions has been reported to promote BAG6-mediated proteasomal degradation—thereby reducing protein abundance (7, 36)—we computed average hydrophobicity for each novel peptide in Kyte–Doolittle [KD] and Miyazawa–Jernigan [MJ] scales using the ‘*peptide*’ R package (37–39). In addition, aggregation properties of the C-terminal peptide sequences were calculated using TANGO and CANYA (40, 41). For TANGO, the maximum score observed among the residues of each C-terminal peptide was selected for analysis. Collectively, the TAILVAR model was built using 37 input features (**Figure 1D**):

**Figure 1.**
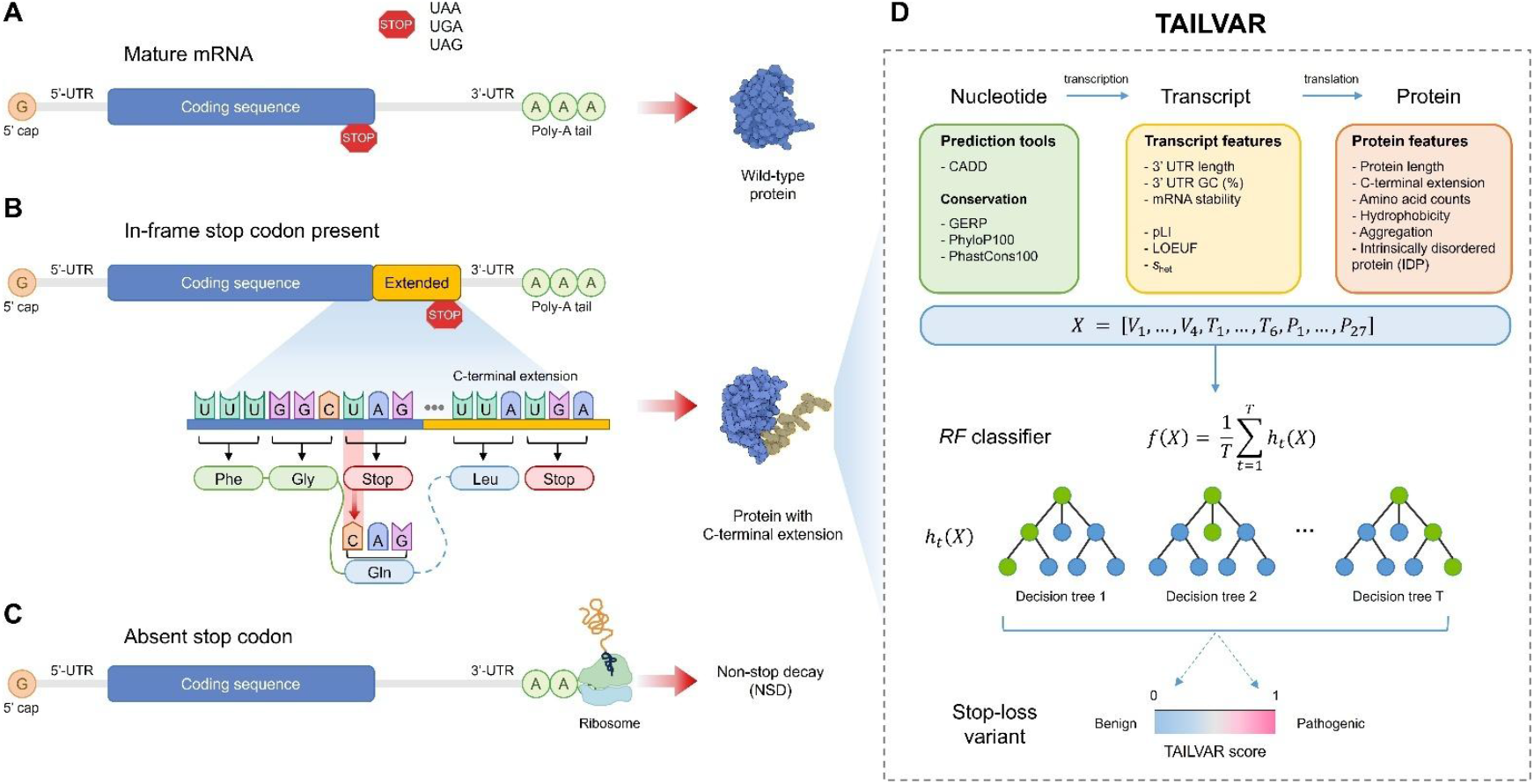
Schematic illustration of translational outcomes by stop-loss variants. (**A**) Structure of a mature mRNA. The mRNA comprises a 5’-untranslated region (UTR), coding sequence, and 3’ UTR, terminating at the stop codon (indicated by “STOP”). Proper translation results in the synthesis of a wild-type protein. (**B**) Impact of stop-loss variants with an in-frame stop codon. Variants at the stop codon convert it to a codon encoding an amino acid, resulting in the continuation of translation and an extension of the open reading frame into the 3’ UTR. If a downstream stop codon is encountered, an aberrant protein containing a C-terminal extension is produced. (**C**) In the absence of a downstream stop codon. If a downstream stop codon is absent, translation proceeds to the end of the mRNA, leading to a stalled ribosome. This can trigger the mRNA surveillance mechanism known as non-stop decay (NSD), resulting in transcript degradation. (**D**) Overview of the TAILVAR model predicting the pathogenicity of stop-loss variants. The TAILVAR model integrates 37 features, encompassing computational prediction scores, transcript, and protein features of the extended portions using random forest (RF) classifiers.

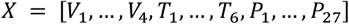

### Model development and assessment

The random forest model (*f*) is an ensemble of decision trees, each trained on a bootstrapped dataset. For the above input data (*X*), the final prediction will be the average of the outputs from all individual trees (*h*_*t*_(*X*)):

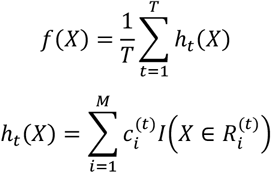

Where *T* is the number of trees (*ntree*), *M* is the maximum number of terminal nodes (*max node*). 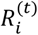is the region associated with *i*th leaf of tree *t*. 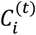 is the constant prediction value for inputs in region 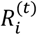. *I*(⋅) is an indicator function, equal to 1 if *X* ∈ 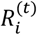, otherwise 0.

Consequently, the model was trained using a 5-fold cross-validation approach to ensure robustness and minimize overfitting. In addition, hyperparameter optimization was conducted through a grid search over the following parameter ranges: *mtry* (number of features randomly selected for splitting at each node: 2, 5, 10, 15, and 20), *ntree* (20, 50, 100, 150, and 200), and *max node* values (5, 10, 20, 30). The optimal hyperparameter combination (*mtry* = 20, *ntree* = 200, *max node* = 30) was selected based on maximizing the area under the receiver operating characteristic (AUROC) curve. Any missing values in the input data were imputed using median imputation. The final model was trained with these optimal hyperparameters and evaluated on both the training and independent test datasets. Model performance was compared to that of 13 established variant effect predictors, including CADD, DANN, Eigen_PC, BayesDel_noAF, BayesDel_addAF, FATHMM_MKL, fitCons_integrated, MutationTaster, VEST4, GPN-MSA, GERP, phyloP100, and phastCons100 (12–22). Performance was assessed using AUROC values calculated with the ‘*pROC*’ package (42). Variable importance was estimated by normalizing the total decrease in the Gini index for each feature. Additionally, a correlation analysis of the input features was performed to evaluate their relative contributions and potential redundancies within the established model.

### Training set

To train the predictive model, we curated a comprehensive dataset of stop-loss variants from large-scale genomic resources. Pathogenic or likely pathogenic (P/LP) variants were extracted from the Human Gene Mutation Database (HGMD Professional, release 2024.2) (43). We selected variant annotated as “DM” (disease mutation) in HGMD, yielding 310 stop-loss variants for the P/LP category. For the benign or likely benign (B/LB) category, we utilized the gnomAD v4.1.0 database, which aggregates 730,947 exomes and 76,215 genomes (30). Stop‐loss variants with a maximum population allele frequency (AF) > 0.1% or observed in the homozygous state were classified as B/LB variants, yielding an additional 309 variants. Variants not located in the MANE transcript or lacking an in-frame stop codon in the 3’ UTR were excluded, resulting in a final training set of 619 stop-loss variants.

### Test set

For model evaluation, we utilized the ClinVar database (release 20250721) and its pathogenicity classifications to categorize variants based on clinical significance (44). Variants classified as ‘Pathogenic’, ‘Likely pathogenic’, or ‘Pathogenic/Likely pathogenic’ were grouped as P/LP (*n* = 187), while those labeled ‘Benign’, ‘Likely benign’, or ‘Benign/Likely benign’ were assigned to the B/LB (*n* = 37) category. Variants with conflicting interpretations of pathogenicity were excluded from analysis. To balance the number of B/LB variants with the P/LP variants, we further incorporated stop-loss variants from multiple genomic resources that provide population AF information. These included Allele Frequency Aggregator (ALFA v2), which aggregates AF information from 192,710 individuals across diverse populations; the Regeneron Million Exome (RME) dataset (45), comprising 983,578 exomes representing varied ancestral backgrounds; and the All Of Us (AOU) Research Program (46), which includes 245,388 genomes. Stop‐loss variants that have an AF greater than 0.1% in any population were regarded as B/LB variants. After removing duplicates, the test set consisted of 558 stop-loss variants.

### Determination of the thresholds

Using a two-component Gaussian mixture model, we established thresholds to distinguish the pathogenicity of variants, as described previously (47). This model was applied to all possible stop-loss variants, utilizing the TAILVAR score as a discriminative parameter. The two components of the model correspond to the probability density distributions for P/LP and B/LB variants. A threshold for the TAILVAR score (*x*_*threshold*_) was established to optimize the identification of potential P/LP and B/LB variants, using a likelihood ratio (LR) below:

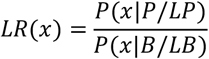

Where *P*(*x*|*P*/*LP*) and *P*(*x*|*P*/*LP*) are the probability density functions (PDFs) for the Gaussian components corresponding to the P/LP and B/LB variant distributions, respectively. These PDFs are defined as:

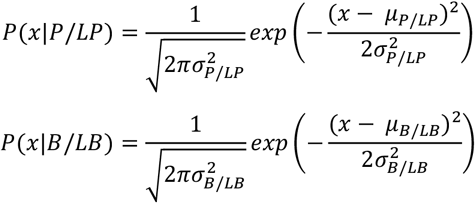

Where μ_*P*/*LP*_ and 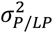 represent the mean and variance of the Gaussian distribution for P/LP variants, and μ_*B*/*LB*_ and 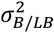 represent the same parameters for B/LB variants. The threshold was determined by solving for the TAILVAR score at which the LR of these two densities equals 9 and 1/9, corresponding to a 90% likelihood of being pathogenic or benign, respectively, using the ‘*mclust*’ R package (**Supplementary Figure S7**) (48). Using these established thresholds, we reassess the pathogenicity of stop-loss variants classified as ‘Uncertain significance’ (VUS) in ClinVar (**Supplementary Table S4**), as well as those in clinically actionable genes listed in the ACMG Secondary Findings (SF) v3.3 genes (**Supplementary Table S5**) (49).

### Correlation analysis with high-throughput functional experiments

To externally validate the robustness of TAILVAR, we analyzed a published dataset comprising high-throughput functional measurements for 2,335 nonstop extension variants identified in cancer (36), cataloged in the NonStopDB (https://NonStopDB.dkfz.de). The NonStopDB variants caused by non-single-nucleotide substitutions were excluded, resulting in 1,848 variants for further evaluation. Functional data for these variants, specifically the median enrichment (ME) scores derived from four biological replicates, were obtained from Supplementary Data 1 of the original publication. These ME scores were log-transformed to normalize the distribution, and subsequent analyses focused on the relationship between these experimental measurements and computational predictions. We compared TAILVAR predictions with 13 additional computational predictions and conservation tools (12–22). For each tool, Spearman correlation coefficients (ρ) were calculated to assess the strength of the relationship between the computational scores and log-transformed ME scores. These correlations were computed using the R function ‘*cor*.*test*’ with the Spearman method, incorporating pairwise complete observations. Additionally, variants were categorized into three TAILVAR score groups (≤ 0.30, 0.30–0.70, ≥ 0.70) based on the established thresholds above. The log-transformed ME scores were compared across these groups to identify trends and validate the discriminative power of TAILVAR.

### Predictions for all possible single-nucleotide substitutions and indels

To construct the TAILVAR database, we systematically pre-computed TAILVAR scores for all possible single-nucleotide substitutions and indels occurring at stop codons across canonical MANE transcripts. For each stop codon, there are three possible single-nucleotide substitutions (SNVs) at each nucleotide position, resulting in a total of nine potential substitutions per codon (see **Table 1**). Similarly, each stop codon permits twelve possible single-nucleotide insertions and three possible single-nucleotide deletions (see **Supplementary Tables S1, S2**). The predicted consequences of the SNVs were annotated using the VEP (28). Due to the issues found in consequence annotation using VEP, indels were annotated using ANNOVAR (50). We excluded variants that resulted in stop-retained alleles or stop-loss alleles lacking an in-frame stop codon within the 3′ UTR. Ultimately, TAILVAR scores were calculated for 306,515 stop-loss variants with C-terminal extensions, comprising 140,334 SNVs, 128,956 insertions, and 37,225 deletions.

**Table 1.** Single-nucleotide substitutions leading to possible codon changes in stop codons

| Stop codon | First nucleotide substitution | Changed amino acids | Second nucleotide substitution | Changed amino acids | Third nucleotide substitution | Changed amino acids |
| --- | --- | --- | --- | --- | --- | --- |
| TGA | (T>A)GA | Arg | T(G>A)A | **Stop<sup>a</sup>** | TG(A>T) | Cys |
|  | (T>G)GA | Gly | T(G>T)A | Leu | TG(A>G) | Trp |
|  | (T>C)GA | Arg | T(G>C)A | Ser | TG(A>C) | Cys |
| TAA | (T>A)AA | Lys | T(A>T)A | Leu | TA(A>T) | Tyr |
|  | (T>G)AA | Glu | T(A>G)A | **Stop<sup>a</sup>** | TA(A>G) | **Stop<sup>a</sup>** |
|  | (T>C)AA | Gln | T(A>C)A | Ser | TA(A>C) | Tyr |
| TAG | (T>A)AG | Lys | T(A>T)G | Leu | TA(G>A) | **Stop<sup>a</sup>** |
|  | (T>G)AG | Glu | T(A>G)G | Trp | TA(G>T) | Tyr |
|  | (T>C)AG | Gln | T(A>C)G | Ser | TA(G>C) | Tyr |

<sup>a</sup>Four substitutions resulting in stop-retained variants are highlighted in bold.

### Statistical analyses

All data used in this work is publicly available (except HGMD, which requires an access request). No statistical method was used to predetermine the sample size. Statistical analyses were performed using the software *R* (v4.4.1). A *P*-value < 0.05 was considered significant throughout the analyses. Categorical variables were compared using a two-tailed Fisher’s exact test, and continuous variables were compared using the Mann⁰Whitney U test. To control the false discovery rate (FDR) for multiple comparisons, the Benjamini–Hochberg procedure was applied, with an FDR threshold of < 0.05.

## Results

### Global impact of stop codon alterations on human transcripts

To investigate the functional impact of variants located in stop codons, we first analyzed 18,890 MANE transcripts. Generally, stop codons define the end point of amino acid translation, with mature mRNA containing poly-A tails downstream in the 3’ UTR (**Figure 1A**). A single nucleotide substitution at a stop codon can yield nine possible outcomes, depending on the specific stop codon involved (**Table 1**). Among these outcomes, stop-retained changes were observed in two out of nine cases for TAA (2/9), while TAG and TGA each have only one stop-retained possibility (1/9). Across all possible SNVs, 14.3% of variants were stop-retained, while 85.7% were stop-loss variants.

For stop-loss variants, translation termination may occur at downstream stop codons within the 3’ UTR (**Figure 1B**) or may continue to the transcript’s end if the stop codon is absent (**Figure 1C**). Transcripts lacking a downstream stop codon are susceptible to NSD, potentially impeding proper protein synthesis. We further examined the predicted consequences of SNVs by assessing in-frame stop codons and alterations in stop codon composition (**Figure 2A**). The distribution of original stop codons across MANE transcripts was TGA (49.2%), TAA (28.4%), and TAG (22.4%). In contrast, downstream stop codon usage shifted to TGA (45.8%), TAA (29.9%), and TAG (20.8%), reflecting an increased presence of TAA. Notably, downstream stop codons were absent in 3.6% of the transcripts, likely susceptible to NSD. Also, we examined the impact of single-nucleotide frameshift indels on stop codon distribution (**Supplementary Tables S1, S2**), and the resulting stop codon frequencies did not differ significantly from those induced by SNVs (**Supplementary Figure S1**).

**Figure 2.**
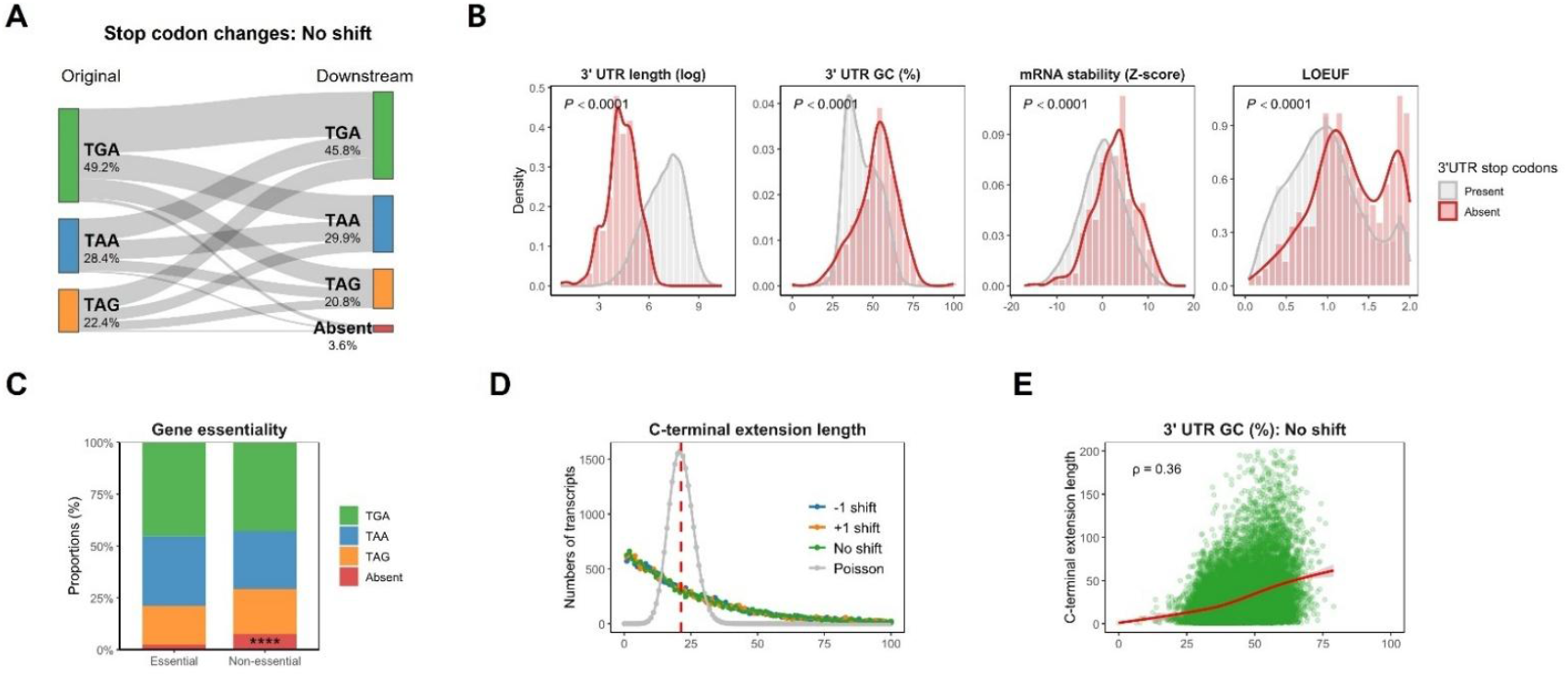
Transcriptome-wide characterization of stop codon usage and C-terminal extension. (**A**) Sankey diagram illustrating transitions in stop codon usage due to single-nucleotide variants (SNVs) that result in stop-loss events across 18,890 MANE transcripts. TGA is the predominant native stop codon, followed by TAA and TAG. Upon stop-loss, downstream codon usage shifts slightly, with 3.6% of transcripts lacking any downstream in-frame stop codon, rendering them susceptible to nonstop decay (NSD). (**B**) Transcript features associated with the absence of downstream stop codons. These transcripts show significantly shorter 3′ UTRs, higher GC content in the 3′ UTR, increased mRNA stability, and elevated LOEUF scores (reduced loss-of-function constraint). (**C**) Proportional usage of downstream stop codons stratified by gene essentiality. Transcripts lacking in-frame stop codons are markedly enriched among non-essential genes, suggesting tolerance to NSD. (**D**) Distribution of C-terminal extension lengths from the original stop to the first downstream in-frame stop codon. Observed distributions deviate from a *Poisson* expectation, supporting a non-random codon structure in the 3′ UTR and potential selection against long, unregulated extensions. Similar distributions are observed following -1 and +1 frameshift events. (**E**) C-terminal extension length had a positive correlation with 3′ UTR GC content (Spearman’s ρ = 0.36). These are consistent with the low GC content of stop codons, which reduces the chance of encountering them.

Next, we examined the distinct characteristics between transcripts with and without downstream in-frame stop codons. As expected, transcripts lacking downstream stop codons had shorter 3’ UTR lengths and higher GC content than those presenting downstream stop codons (**Figure 2B**). Additionally, transcripts without downstream stop codons exhibited increased mRNA stability, elevated LOEUF scores, and enrichment in non-essential genes (**Figure 2C**). The distinct transcript features in 3’ UTR lengths, mRNA stability, and LOEUF scores were also noted by gene essentiality classifications (**Supplementary Figure S2**). These findings suggest that the presence of downstream stop codons is potentially under selective pressure. Additionally, we examined the distribution of C-terminal extension lengths across all human transcripts (**Figure 2D**). Under a random nucleotide model, the expected extension length follows a *Poisson* distribution with a mean of 21.3 (64/3) codons, assuming uniform codon usage. However, the observed distribution was significantly skewed toward shorter extensions, with a median of 19 amino acids, suggesting that 3′ UTRs harbor sequence constraints that limit extension length. Remarkably, frameshift events introduced by +1 shift or -1 shift produced similar extension length distributions, indicating that selective pressures impact broadly across all reading frames (**Supplementary Figure S1**). Extension length also exhibited a moderate positive correlation with 3′ UTR GC content (Spearman’s ρ = 0.36; **Figure 2E**), consistent with the known AT-rich composition of stop codons. As stop codons (TAA, TAG, TGA) are enriched for A and T nucleotides, GC-rich regions are inherently depleted of termination signals, increasing the likelihood of extended translation.

### Characterization of stop-loss variants contributing to pathogenicity

To assess the biological features that distinguish B/LB from P/LP stop-loss variants, we analyzed the variant-, transcript-, and protein-level metrics across our curated datasets (**Figure 3A**): the training set (HGMD/gnomAD; *n* = 619) and the test set (ClinVar/ALFA/RME/AOU; *n* = 558). Interestingly, we observed that 3’ UTR GC contents, length of 3’ UTR, mRNA stability, and length of the C-terminal extension were greater in the P/LP classification, while higher LOEUF values were noted in the B/LB classification, reflecting lower selective constraints (**Figure 3B**).

**Figure 3.**
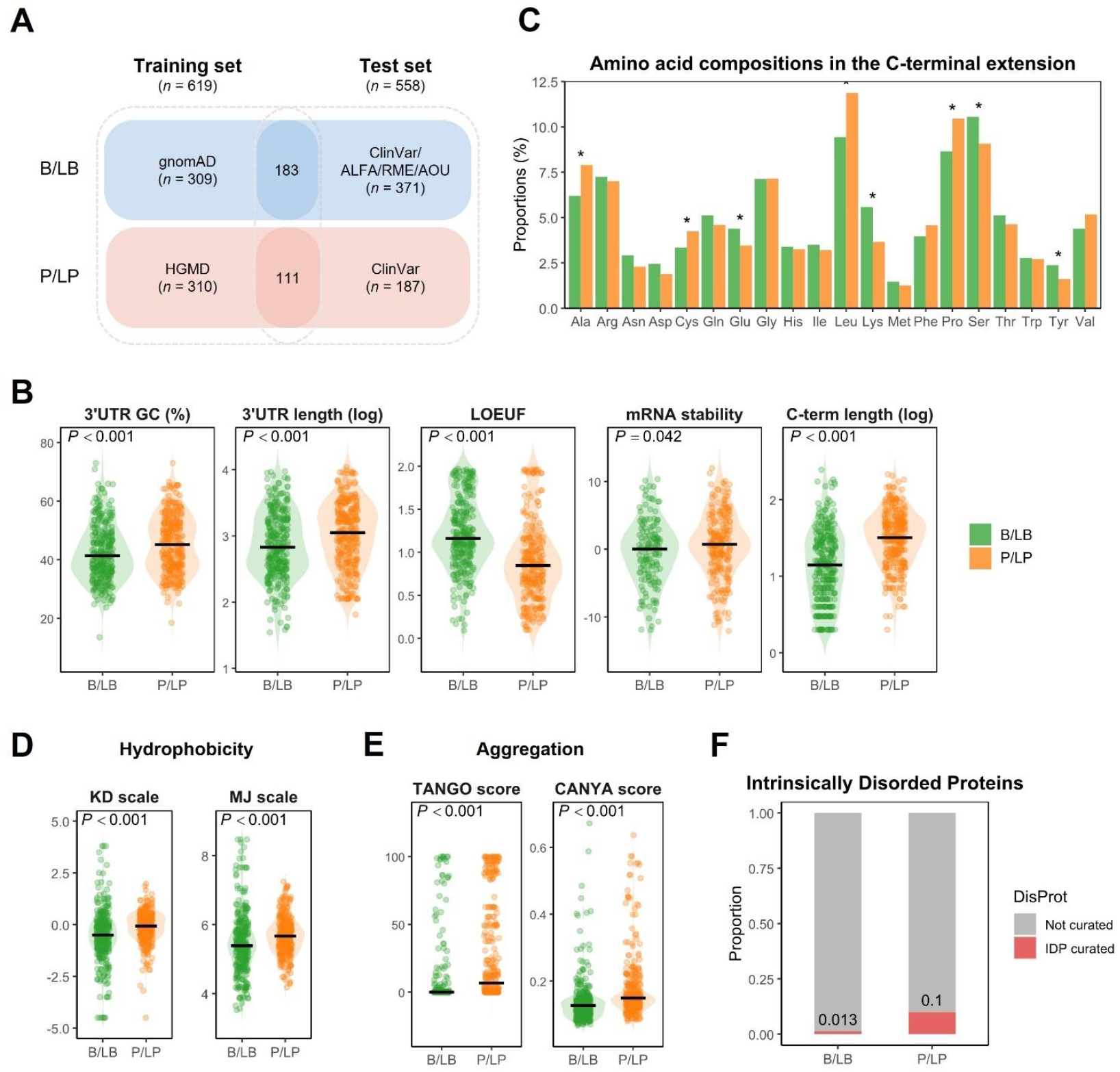
Multi-omics features influence the functional impact of elongated proteins. (**A**) Overview of the curated datasets. The training set comprises stop-loss variants classified as benign/likely benign (B/LB; blue) from gnomAD and pathogenic/likely pathogenic (P/LP; red) from HGMD, while the test set includes B/LB variants from ClinVar, ALFA, RME, or AOU and P/LP variants from ClinVar. (**B**) Violin plots comparing transcript-level features between B/LB (green) and P/LP (orange) variants in the training set. From left to right: 3′ UTR GC content (percentage), 3′ UTR length (log-transformed), LOEUF (loss-of-function observed/expected upper bound), mRNA stability (Z-score), and C-terminal extension length (log-transformed). *P*-values for group comparisons are indicated above each plot. (**C**) Amino acid composition of C-terminal extensions for B/LB and P/LP variants. Residues significantly enriched in P/LP extensions (Ala, Cys, Leu, Pro) and those enriched in B/LB extensions (Glu, Lys, Ser, Tyr) are marked with asterisks (adjusted *P* < 0.05). (**D**) Hydrophobicity comparison of C‐terminal extensions using the Kyte–Doolittle (KD) and Miyazawa–Jernigan (MJ) scales. Violin plots show that P/LP‐derived peptides have significantly higher average hydrophobicity than B/LB peptides. (**E**) Aggregation propensity of C-terminal extensions as measured by TANGO and CANYA scores. P/LP variants exhibit significantly higher aggregation scores compared to B/LB. (**F**) Proportion of intrinsically disordered proteins (IDPs) among genes harboring stop‐loss variants, based on DisProt annotation. P/LP variants (right) show a higher IDP fraction compared to B/LB variants.

In addition, we examined amino acid usage at the original stop codon as well as in the aberrant C‐terminal extensions. Although no single stop‐codon–to–amino‐acid change reached statistical significance, arginine (Arg) was the most frequently observed amino acid at stop codon in P/LP variants, whereas glutamine (Gln) was the most frequently observed amino acid at stop codon in B/LB variants (**Supplementary Figure S3**). In the C‐terminal extensions themselves, P/LP variants were enriched for alanine (Ala), cysteine (Cys), leucine (Leu), and proline (Pro)—residues associated with hydrophobicity and conformational rigidity—whereas B/LB extensions contained more glutamate (Glu), lysine (Lys), serine (Ser), and tyrosine (Tyr) which are hydrophilic or flexible (**Figure 3C**). Consistent with these observations, average hydrophobicity scores (Kyte–Doolittle [KD] and Miyazawa–Jernigan [MJ] scales) were significantly higher in P/LP‐derived C-terminal peptides (**Figure 3D**). Furthermore, aggregation propensities, measured by TANGO and CANYA scores, were significantly higher in the C-terminal extensions of P/LP variants, highlighting distinct protein features (**Figure 3E**). Finally, we investigated the proprotions of curated IDPs using the DisProt database. Remarkably, IDPs were significantly more enriched in the P/LP variants (10.0%) than B/LB variants (1.3%, **Figure 3F**). All of these observations in the training set were consistently replicated in the test set (**Supplementary Figure S4**). Collectively, these findings indicate that pathogenic events preferentially occur in genes with disordered C-terminal regions, producing longer, aggregation-prone, hydrophobic extensions, and suggest a mechanism by which disruption of IDP-rich tails may contribute to disease.

### Development and assessment of TAILVAR

Based on the observations above, we developed a machine-learning classifier, TAILVAR (Terminal codon Analysis and Improved prediction of Lengthened VARiants), to accurately predict the functional consequences of stop-loss variants (**Figure 1D**). Notably, the total number of variants in the training and test datasets was limited, reflecting the rarity of stop-loss variants. Indeed, these variants represent only 0.04% of all recorded entries in the ClinVar database (**Supplementary Table S3**). For model development, we incorporated 37 features, encompassing computational prediction (CADD), conservation scores, and transcript and protein properties of the C-terminal extensions analyzed above (**Figure 4A**). CADD and conservation scores exhibited strong intercorrelations, whereas transcript and protein features showed lower correlation with these tools, thereby offering additional predictive value (**Supplementary Figure S6**).

**Figure 4.**
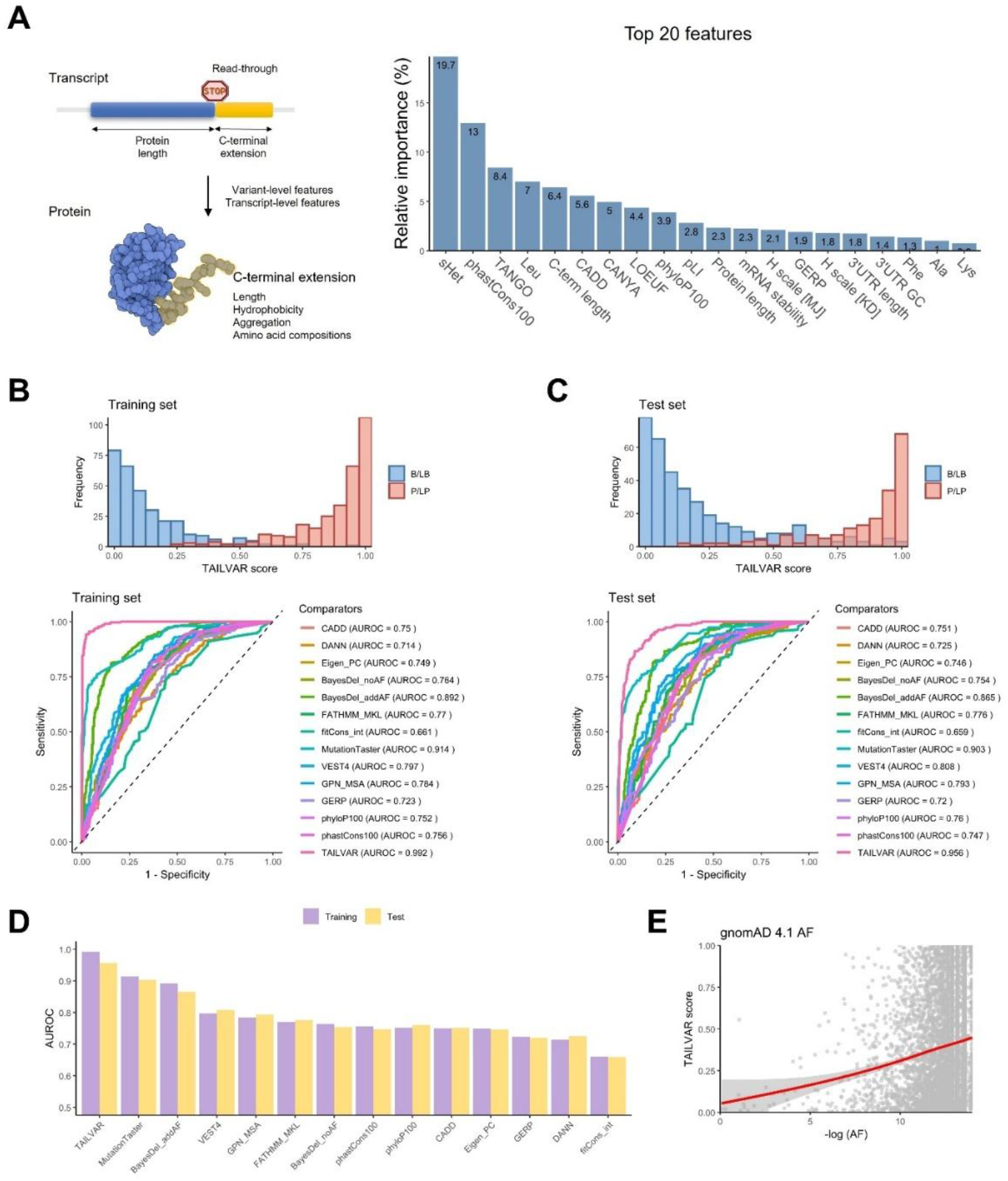
Overview and benchmark results of TAILVAR. (**A**) Schematic overview of the TAILVAR framework, highlighting the transcript and protein-level features incorporated into model development (left), alongside the top 20 most important features ranked by normalized Gini index (right). Among these, phastCons100, TANGO, CADD, and leucine (Leu) content are among the top contributors to model performance. (**B**) Distribution of TAILVAR scores for benign/likely benign (B/LB; blue) and pathogenic/likely pathogenic (P/LP; red) variants in the training set, with corresponding receiver operating characteristic (ROC) curves demonstrating discrimination capacity across TAILVAR and 13 comparator prediction tools. TAILVAR achieves a superior AUROC (0.992) in the training set. (**C**) Distribution of TAILVAR scores and ROC curves in the independent test set, with P/LP and B/LB variants effectively separated. TAILVAR maintains the highest AUROC (0.956) among all tools assessed. (**D**) Comparative barplot of AUROC values for TAILVAR and other benchmark tools in the training (purple) and test (yellow) datasets, confirming consistently high performance of TAILVAR across datasets. (**E**) Scatterplot showing the relationship between TAILVAR scores and allele frequency (AF) for variants in gnomAD v4.1. A negative correlation is observed, indicating that variants predicted as more pathogenic by TAILVAR tend to have lower population frequencies.

Among the features included, constraint and conservation metrics—such as *s*_het_ and phastCons100—ranked first and second, respectively, in importance, as measured by the decrease in the Gini index, highlighting their strong predictive performance. Notably, characteristics of the C-terminal extensions, including aggregation scores (TANGO and CANYA), C-terminal length, and hydrophobicity (with leucine as a representative hydrophobic amino acid), also contributed significantly, emphasizing the critical role of protein-specific properties. Overall, these findings support that increased C-terminal extension length and hydrophobicity likely enhance susceptibility to ubiquitin-mediated proteasomal degradation, whereas elevated aggregation propensity may promote the formation of toxic mutant proteins, thereby amplifying pathogenic potential (36, 41).

The TAILVAR model demonstrated remarkable discrimination between P/LP and B/LB variants, with an AUROC of 0.992 in the training set and outperforming all other benchmarks (**Figure 4B**). To assess potential overfitting, we further evaluated TAILVAR on an independent test set, where it maintained the highest performance with an AUROC of 0.956, confirming its robust predictive capacity (**Figure 4C**). Among the alternative tools assessed, MutationTaster and BayesDel_addAF ranked second and third, respectively—likely reflecting their incorporation of AF information (**Figure 4D**). Notably, although TAILVAR does not utilize AF as an input, analysis of its predicted scores for gnomAD variants revealed a negative correlation with AF, indicating that variants with higher predicted pathogenicity tend to be rarer in the population (**Figure 4E**).

### Application of TAILVAR in human genetics

To establish robust thresholds for the TAILVAR score, we applied a two-component Gaussian mixture model to all possible stop-loss variants (**Figure 5A**). Thresholds were set at 0.30 and 0.70, corresponding to a 90% probability of correct classification as benign or pathogenic, respectively (**Supplementary Figure S7**). Using these criteria, 50,772 (36.2%) variants were classified as potentially P/LP, and 47,835 (34.1%) as B/LB. Applying the same thresholds (TAILVAR score ≥ 0.70 and ≤ 0.30) to VUS variants in the ClinVar database, 287 (41.7%) and 173 (25.1%) VUS variants in the ClinVar database could be reclassified as VUS-P (potentially P/LP) and VUS-B (potentially B/LB), respectively (**Figure 5B**; **Supplementary Table S4**). Furthermore, we identified 359 potential P/LP variants across 84 clinically actionable ACMG genes, highlighting their clinical significance (**Supplementary Table S5**) (49). Notably, the majority of these variants had not been previously reported, further emphasizing the utility of TAILVAR in identifying novel pathogenic candidates.

**Figure 5.**
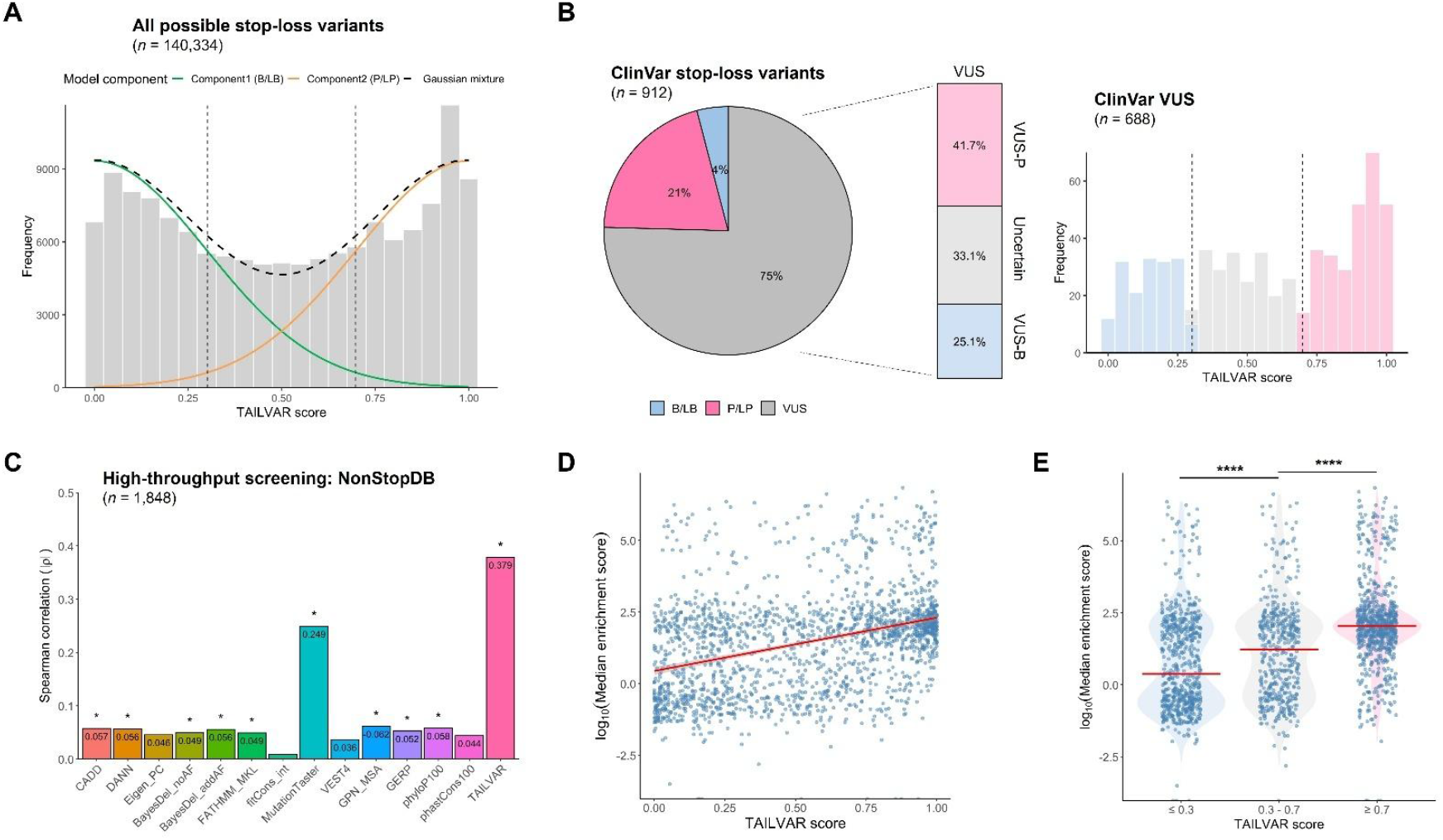
TAILVAR facilitates reclassification of clinically significant stop-loss variants and correlates with functional experiments. (**A**) Two-component Gaussian mixture modeling of TAILVAR scores for all possible stop-loss variants (*n* = 140,334), delineating distributions corresponding to benign/likely benign (B/LB) and pathogenic/likely pathogenic (P/LP) classes. Dashed lines indicate thresholds (0.30 and 0.70) used for variant classification. (**B**) Distribution of ClinVar stop-loss variants (*n* = 912) categorized as B/LB, P/LP, and variants of uncertain significance (VUS). The right panels show further stratification of VUS variants (*n* = 688) using TAILVAR thresholds into potentially pathogenic (VUS-P; ≥ 0.70), uncertain (0.30–0.70), and potentially benign (VUS-B; ≤ 0.30) groups, along with their respective proportions. (**C**) Correlation analysis between computational prediction scores and high-throughput screening data for cancer-derived nonstop extension variants (NonStopDB) in HEK293T cells (*n* = 1,848). Spearman correlation coefficients (ρ) are shown for TAILVAR and 13 additional computational tools, with significance indicated by asterisks (\**P* < 0.05). TAILVAR demonstrates the strongest correlation (ρ = 0.379) with experimental measurements. (**D**) Scatter plot between TAILVAR scores and log-transformed median enrichment score derived from the high-throughput functional screening data. The observed positive correlation indicates that higher TAILVAR scores are associated with decreased protein abundance. (**E**) Violin plots comparing log-transformed median enrichment scores across TAILVAR score categories (≤ 0.30, 0.30–0.70, ≥ 0.70), showing significant differences among groups (\*\*\*\**P* < 0.0001).

The peptide sequences and lengths of the C-terminal extensions were found to depend primarily on the transcript, regardless of the variant position, except for the specific amino acid substituted at the original stop codon. Because the resulting protein generally preserves its amino acid sequence aside from this single amino acid change, the pathogenicity of stop-loss variants arising from the same transcript is likely to be similar. Leveraging these insights and TAILVAR scores, we were able to reclassify 31 VUS-P variants across 25 genes as P/LP variants (**Supplementary Table S6**). Among the 25 genes, we identified six genes with additional evidence supporting the pathogenicity of VUS through a comprehensive literature review (**Table 2**). For instance, in *GCH1* (ENST00000491895), which is associated with Dopa-responsive dystonia (MIM #600225), two variants (c.751T>C, p.Ter251ArgextTer35; c.752G>C, p.Ter251SerextTer35) were classified as P, and two other variants (c.753A>C, p.Ter251CysextTer35; c.753A>G, p.Ter251TrpextTer35) were classified as VUS in ClinVar. These variants have identical peptide sequences except for the amino acids at the original stop codon (Arg, Ser, Cys, and Trp, respectively). Given that the only difference is this single amino acid substitution, and considering the high TAILVAR scores (1.000) for the VUS variants, p.Ter251CysextTer35 and p.Ter251TrpextTer35 warrant reclassification as P/LP. Indeed, this interpretation is supported by a prior clinical report documenting a large Danish family with Dopa-responsive dystonia carrying the p.Ter251CysextTer35 variant (51). Similarly, in *ARSB* (ENST00000264914), which is associated with mucopolysaccharidosis type VI (Maroteaux-Lamy syndrome; MIM #253200), the VUS variant p.Ter534GlnextTer50 showed a high TAILVAR score (0.995), as did two reported P/LP variants (with Ser and Trp at the stop codon site) within the same transcript. Functional studies demonstrated that the p.Ter534GlnextTer50 variant is rapidly degraded before reaching the trans-Golgi, providing molecular evidence for its pathogenicity (52).

**Table 2.**
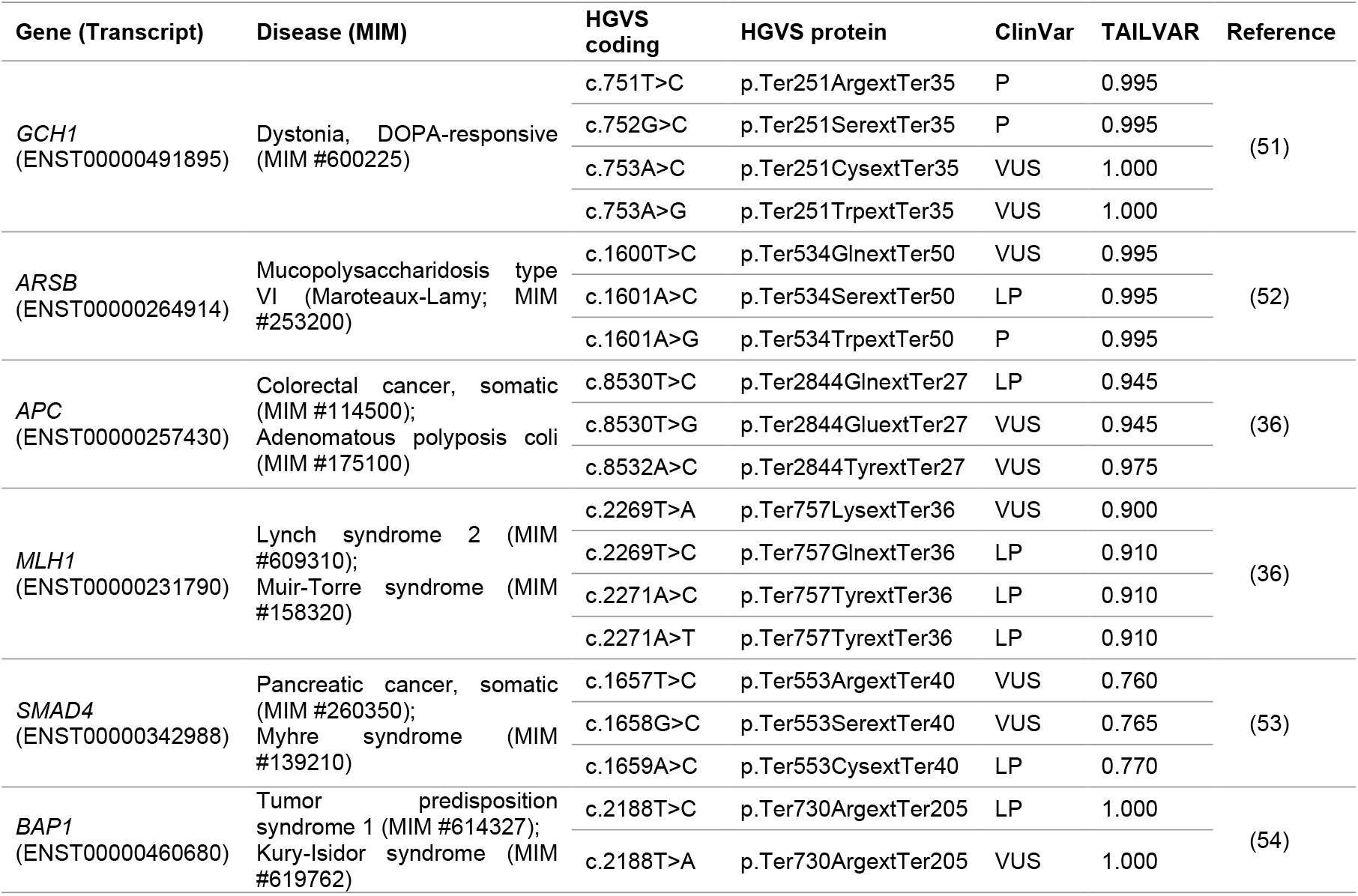
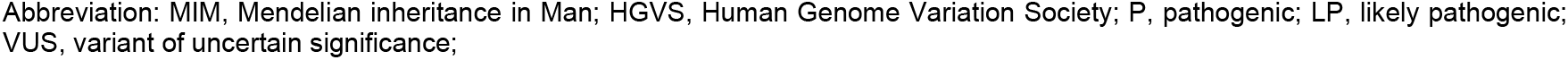
Reclassification of VUS as P/LP supported by pathogenic stop-loss variants with identical C-terminal extensions and literature evidence

| Gene (Transcript) | Disease (MIM) | HGVS coding | HGVS protein | ClinVar | TAILVAR | Reference |
| --- | --- | --- | --- | --- | --- | --- |
| *GCH1*<br>(ENST00000491895) | Dystonia, DOPA-responsive<br>(MIM #600225) | c.751T>C | p.Ter251ArgextTer35 | P | 0.995 | (51) |
|  |  | c.752G>C | p.Ter251SerextTer35 | P | 0.995 |  |
|  |  | c.753A>C | p.Ter251CysextTer35 | VUS | 1.000 |  |
|  |  | c.753A>G | p.Ter251TrpextTer35 | VUS | 1.000 |  |
| *ARSB*<br>(ENST00000264914) | Mucopolysaccharidosis type VI (Maroteaux-Lamy; MIM #253200) | c.1600T>C | p.Ter534GlnextTer50 | VUS | 0.995 | (52) |
|  |  | c.1601A>C | p.Ter534SerextTer50 | LP | 0.995 |  |
|  |  | c.1601A>G | p.Ter534TrpextTer50 | P | 0.995 |  |
| *APC*<br>(ENST00000257430) | Colorectal cancer, somatic (MIM #114500);<br>Adenomatous polyposis coli (MIM #175100) | c.8530T>C | p.Ter2844GlnextTer27 | LP | 0.945 | (36) |
|  |  | c.8530T>G | p.Ter2844GluextTer27 | VUS | 0.945 |  |
|  |  | c.8532A>C | p.Ter2844TyrextTer27 | VUS | 0.975 |  |
| *MLH1*<br>(ENST00000231790) | Lynch syndrome 2 (MIM #609310);<br>Muir-Torre syndrome (MIM #158320) | c.2269T>A | p.Ter757LysextTer36 | VUS | 0.900 | (36) |
|  |  | c.2269T>C | p.Ter757GlnextTer36 | LP | 0.910 |  |
|  |  | c.2271A>C | p.Ter757TyrextTer36 | LP | 0.910 |  |
|  |  | c.2271A>T | p.Ter757TyrextTer36 | LP | 0.910 |  |
| *SMAD4*<br>(ENST00000342988) | Pancreatic cancer, somatic (MIM #260350);<br>Myhre syndrome (MIM #139210) | c.1657T>C | p.Ter553ArgextTer40 | VUS | 0.760 | (53) |
|  |  | c.1658G>C | p.Ter553SerextTer40 | VUS | 0.765 |  |
|  |  | c.1659A>C | p.Ter553CysextTer40 | LP | 0.770 |  |
| *BAP1*<br>(ENST00000460680) | Tumor predisposition syndrome 1 (MIM #614327);<br>Kury-Isidor syndrome (MIM #619762) | c.2188T>C | p.Ter730ArgextTer205 | LP | 1.000 | (54) |
|  |  | c.2188T>A | p.Ter730ArgextTer205 | VUS | 1.000 |  |

These findings also extend to cancer predisposition genes, such as *APC* (ENST00000257430), *MLH1* (ENST00000231790), *SMAD4* (ENST00000342988), and *BAP1* (ENST00000460680). A previous study demonstrated that nonstop extensions in *APC* (p.Ter2844GluextTer27) and *MLH1* (p.Ter757LysextTer36, p.Ter757TyrextTer36) indeed result in a loss of protein expression (36). Furthermore, *SMAD4* stop codon substitutions to Arg and Ser (p.Ter553ArgextTer40 and p.Ter553SerextTer40) were shown to be degraded by the ubiquitin-mediated proteasomal pathway (53). TAILVAR scores were consistent with these functional experimental results and with previously reported LP variants in the same transcripts. Similarly, mechanistic studies of *BAP1* stop-loss variants (Arg, Gly, Leu, and Ser) have revealed reduced protein translation via a degradation-independent mechanism (54). Notably, the *BAP1* c.2188T>C and c.2188T>A variants both produce an identical protein product with Arg at the original stop codon (p.Ter730ArgextTer205). Thus, the identical protein consequence justifies assigning the same clinical classification to both variants, with the VUS warranting reclassification as LP.

Deep mutational scanning (DMS) or multiplexed assays of variant effect (MAVE) are widely used for the functional evaluation of missense variants (23). The correlation of *in silico* prediction scores with these datasets is an important metric for assessing their biological relevance. For this purpose, we utilized a publicly available dataset containing high-throughput functional screening data of cancer-derived nonstop C-terminal extensions (NonStopDB) (36). This dataset measured protein abundance of nonstop extensions using fluorescence-activated cell sorting with four replicates in HEK293T cells. A comparison of Spearman correlation coefficients (ρ) between TAILVAR and 13 benchmark scores indicates that TAILVAR achieved the highest correlation (ρ = 0.379) with this experimental data, surpassing other tools, which showed weak or very weak correlations (**Figure 5C**). Although this correlation is moderate, it is notable given the inherent variability of the assay—biological replicates in the original study exhibited ρ values ranging from 0.65 to 0.86. Using the established score thresholds, TAILVAR also distinguished variants with significantly different protein abundance levels, demonstrating both strong discriminatory power and biological relevance (**Figure 5D, E**). Importantly, the high-throughput assay measures protein abundance and is therefore primarily suited to detecting LOF effects. Variants with GOF consequences, which may alter activity without changing abundance, are less likely to be captured by this approach, potentially contributing to the relatively modest correlations. Overall, these results indicate that TAILVAR correlates well with experimental measurements and reliably reflects the protein abundance of nonstop C-terminal extensions.

Finally, we conducted an additional literature review and identified 17 genes harboring 36 stop-loss variants characterized by well-established functional experiments, including *ITM2B, FOXE3, MITF, FLNA, VHL*, and *PTEN* (**Supplementary Table S7**) (36, 54–58). Many of these variants are cataloged in the HGMD database; 16 were included in the TAILVAR training set and exhibited high scores. Notably, 20 variants not present in the training set—including those in *APC, BAP1, PTEN, SMAD4*, and *FLNA*—also received high TAILVAR scores, further supporting the model’s reliability and its concordance with experimental data reported in the literature. Most stop-loss variants exert a LOF effect, primarily via C-terminal extensions that promote proteasomal degradation. However, some genes, including *APOA2, FGFR3, ITM2B*, and *FLNA*, have been associated with gain-of-function (GOF) effects (57–60). Taken together, these findings highlight the strong agreement between TAILVAR predictions and experimental or literature-based evidence, while also emphasizing that elucidating the precise disease mechanisms—whether LOF or GOF— remains an important area for future investigation.

### Interpretive framework for stop-loss variants

Although stop-loss variants represent a relatively small subset of genomic variants, their pathogenicity is occasionally underestimated in clinical assessments (**Supplementary Table S3**). This underestimation arises mainly from limited knowledge and the absence of a robust interpretive framework. In general, stop-loss variants that result in C-terminal extensions are classified as having moderate pathogenic evidence (PM4) according to the ACMG/AMP criteria (10). Recently, the Association for Clinical Genomic Science (ACGS) Best Practice Guidelines have refined this framework, proposing PVS1 for stop-loss variants lacking an in-frame stop codon in the 3’ UTR, which can lead to NSD (61). Based on our analyses, we propose additional considerations and present a comprehensive workflow to enhance the interpretation of stop-loss variants (**Figure 6**).

**Figure 6.**
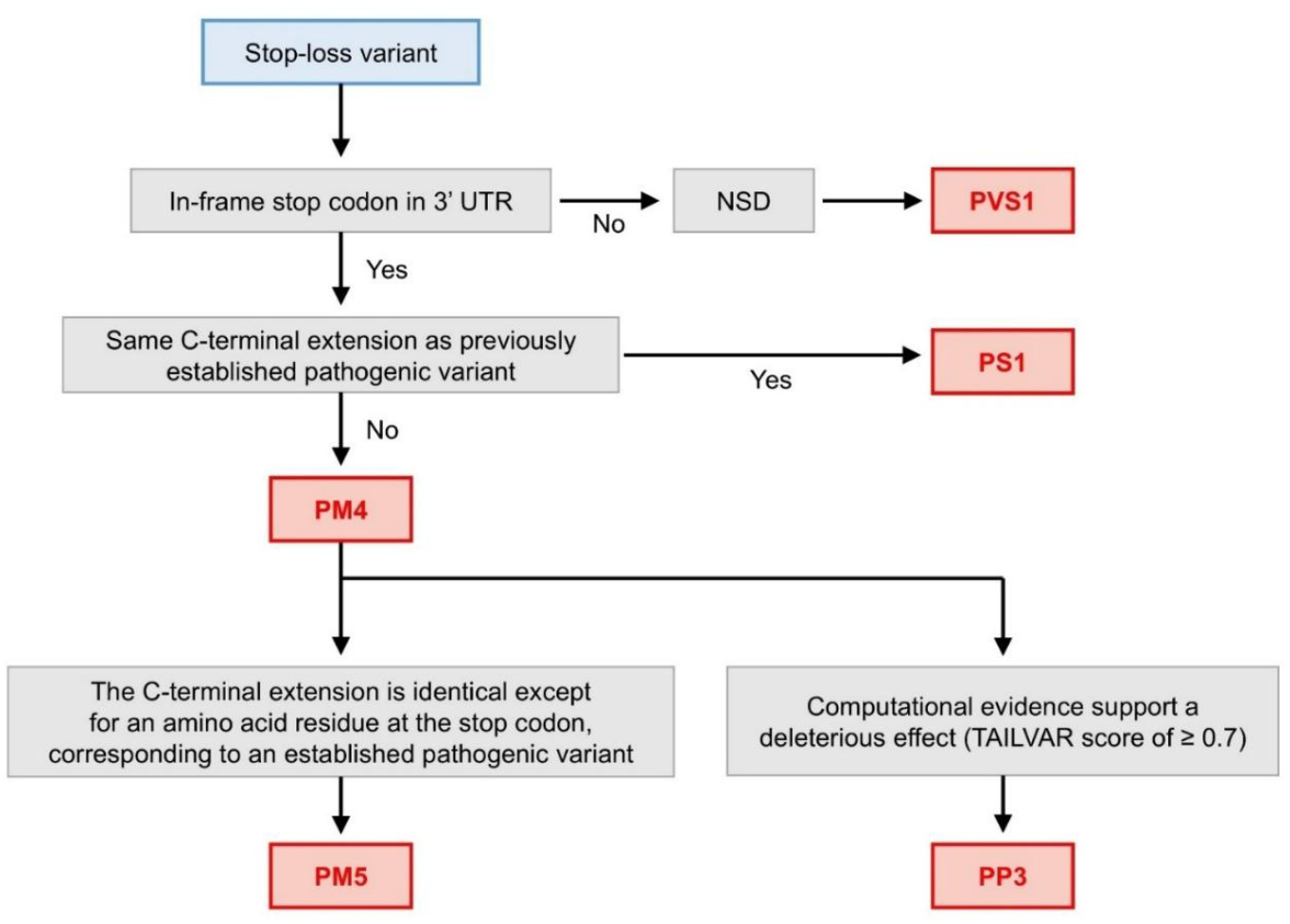
Proposed flowchart for assessing the pathogenicity of stop-loss variants. This flowchart outlines a systematic approach for evaluating stop-loss variants, integrating the established ACMG/AMP criteria. Variants are first assessed for the presence of an in-frame stop codon within the 3’ untranslated region (3’ UTR). If no downstream stop codon exists, non-stop decay (NSD) leads to protein degradation, qualifying the variant for the PVS1 criterion. Conversely, if an in-frame stop codon is present, the analysis examines whether the resulting C-terminal extension matches that of a previously established pathogenic variant (PS1). If not, it can be categorized into the PM4 criterion (protein length changes), the same as the ACMG/AMP criteria. However, additional considerations apply if the C-terminal extension is identical to that of an established pathogenic variant except for the amino acid at the stop codon (PM5), or if there is supporting computational evidence for pathogenicity, such as a TAILVAR score ≥ 0.7 (PP3). This framework provides a stepwise method to refine the classification of stop-loss variants based on their protein features and functional impacts.

As highlighted in the ACGS guidelines, the absence of an in-frame stop codon is a critical predictor of NSD and provides a valuable starting point for interpreting stop-loss variants. Our analysis reveals that approximately 3.6% of transcripts with single-nucleotide substitutions or indels fall into this category (**Figure 2A, Supplementary Figure S1**). In addition to NSD prediction, further refinement of stop-loss variant interpretation can be achieved by benchmarking against missense variant criteria (**Supplementary Table S8**). For example, analogous to the PS1 criterion for missense variants, stop-loss variants predicted to produce identical C-terminal extensions with established pathogenicity should be evaluated similarly. In our analysis, two *BAP1* stop-loss variants, c.2188T>C and c.2188T>A, both result in the identical extension p.Ter730ArgextTer205 (**Table 2**). While one is classified as LP and the other as VUS in ClinVar, we suggest that the VUS should also be classified as LP (PS1, PM2).

Additionally, our analysis of stop-loss variants caused by single-nucleotide substitutions revealed that the resulting elongated proteins differ only at the amino acid corresponding to the original stop codon, as seen in *GCH1, ARSB, APC, MLH1*, and *SMAD4* (**Table 2**). Variants that affect the same elongated protein can therefore be regarded as nonsynonymous variants of the extended protein. Accordingly, if a nonstop extension has been previously established as pathogenic, moderate evidence of pathogenicity (PM5 criterion) can be assigned to a variant that differs solely at the stop codon amino acid, particularly when supported by similar TAILVAR scores. In addition, a TAILVAR score ≥ 0.7 can be incorporated into the PP3 criterion, providing further computational evidence for pathogenicity. Overall, our proposed framework (**Figure 6**) offers a systematic approach to refining stop-loss variant classification. By integrating factors such as NSD prediction, similarity to C-terminal extensions with established pathogenicity, and quantitative assessment of the extension using a machine learning classifier, this workflow addresses current gaps and enables more accurate and efficient interpretation of stop-loss variants.

## Discussion

To our knowledge, no computational tools have been specifically developed for stop-loss variants, largely owing to their rarity. Yet, these variants are recurrently linked to pathogenic outcomes, highlighting the critical need for dedicated methods to assess their clinical impact. In the expanding era of genetics and genomics, accurate prediction and interpretation of all variant types are essential for advancing precision medicine. This includes specialized approaches for underrepresented classes such as start-loss and stop-loss variants. As part of this effort, the PoStaL model has provided valuable insights into start-loss variants (11). Herein, we introduce TAILVAR—a model designed to address the functional impacts of C-terminal extension caused by stop-loss variants. By integrating established predictive frameworks with transcript- and protein-level context, TAILVAR offers a systematic and comprehensive approach to interpreting this distinctive variant class (**Figure 1D**).

The transcriptome-wide analysis of stop codon usage and shifts induced by stop-loss variants has uncovered several notable insights (**Figure 2A**). We found that the patterns of stop codon usage observed in coding regions are largely maintained in downstream stop codons, with a slightly increased usage of TAA. Among the three canonical stop codons, TAA is considered the most optimal and robust, exhibiting the lowest rates of translational readthrough (62). Consistent with this, TAA demonstrated a higher probability of being retained as a stop codon following mutations compared to TAG and TGA, demonstrating its stability (**Table 1, Supplementary Tables S1, S2**). This increased shift to TAA may also be influenced by the presence of polyadenylation signals, particularly the frequent AAUAAA motifs within the 3′ UTR (63). In contrast, TGA emerged as the most frequently used stop codon in the human genome, while TAG was the least utilized—a phenomenon often referred to as the TAG-TGA paradox. Recent studies have attributed this paradox to methylation and the hypermutability of CpG dinucleotides, which preferentially lead to TGA codons (64). Notably, approximately 3.6% of transcripts were found to lack any in-frame stop codons downstream. These transcripts are characterized by shorter 3′ UTR lengths, higher 3′ UTR GC content relative to others, and lower selective constraint (**Figure 2B**). This observation aligns with prior findings suggesting that stop codon usage in the human genome is shaped by both GC content in coding regions and gene expression levels. For example, highly expressed genes tend to preferentially favor AT-rich sequences (65), which are correlated with shorter C-terminal extensions (**Figure 2E**) and may mitigate the potential toxic effects of longer C-terminal extensions. On the other hand, highly expressed genes are also more likely to develop intrinsically disordered C-termini, which can minimize the deleterious effects of C-terminal extensions without compromising native protein function (5). Furthermore, the peptide sequences of highly expressed genes introduced by the 3’UTR tend to have a higher fraction of hydrophobic amino acids, making them more susceptible to proteasomal degradation (7). Collectively, these mechanisms may help preserve gene function, even as genes accumulate higher GC content through evolutionary selection.

Our analyses reveal distinct transcript and protein signatures that differentiate pathogenic potentials (**Figure 3, Supplementary Figure S4**). Notably, the amino acid composition of C-terminal extensions diverges significantly between P/LP and B/LB groups (**Figure 3C**). P/LP variants are enriched in hydrophobic and conformationally rigid residues, with Leu, Pro, Ala, and Cys following this order of proportions, whereas B/LB variants have significantly higher proportions of hydrophilic and conformationally flexible residues such as Ser, Lys, and Glu. In particular, Leu, known for its strong hydrophobic nature, is occasionally buried in the protein core, where it stabilizes the hydrophobic interior (37). Likewise, Pro has a unique cyclic structure, formed by covalent bonding of its side chain to the nitrogen atom of the amino group, restricting N-Cα bond rotation, making it the most conformationally constrained amino acid (66, 67). In contrast, Ser and Lys are polar and exhibit higher structural tolerance, as their presence is less likely to disrupt the protein core or essential interactions (68). Together, the average hydrophobicity of the C-terminal extension was remarkably different between P/LP and B/LB variants (**Figure 3D**). These findings align with previous observations that increased hydrophobicity in C-terminal extensions can induce proteasomal degradation via the BAG6 pathway, thereby preventing the accumulation of non-canonical proteins (7). The higher hydrophobicities are likely to induce proteasomal degradation, resulting in P/LP variants as demonstrated in *SMAD4* nonstop extensions (53).

Another key feature of pathogenic C-terminal extensions is their propensity for aggregation (**Figure 3D**). The formation of protein aggregates is well-established as a source of toxic GOF, as observed in neurodegenerative disorders such as Parkinson’s disease, amyotrophic lateral sclerosis, and familial British dementia associated with *ITM2B* (57), as well as in systemic amyloidosis (60). Aggregation can also contribute to LOF phenotypes by impairing or altering the normal activity of proteins, and may elicit unfolded protein responses that lead to endoplasmic reticulum (ER) stress (69). Thus, the high predictive value of TANGO and CANYA scores in the TAILVAR model is mechanistically plausible (**Figure 4A**). Additionally, neopeptide sequences introduced by C-terminal extensions may encode novel cellular localization signals, which may alter their functions and interaction with other molecules. While our current model did not include this feature due to the limited number of such cases, future models could be further enhanced by incorporating the information on altered subcellular localization (70).

Despite the high predictive accuracy, our model has several limitations. First, the number of variants used for training and evaluation is relatively small, constraining its applicability to advanced machine learning techniques such as deep learning. Adopting similar approaches to those employed for missense variants—such as curating larger variant collections or incorporating synthetic data through DMS or MAVE—could address this limitation in future iterations (71). Nevertheless, our study demonstrated the highest correlation with the high-throughput screening data of nonstop extension mutations (**Figure 5C**) (36). Incorporating larger experimental datasets in the future would significantly enhance the predictive power of stop-loss variants. Second, the model assumes that the nearest downstream stop codon in the 3’ UTR is always utilized, an assumption that may not hold for all variants. Stop codon readthrough or changes in mRNA secondary and tertiary structure could alter stop codon selection (8, 72), adding complexity that is not currently captured. Incorporating mRNA 3D structural change predictions into future iterations of the model may enhance its predictive accuracy (73). Third, by focusing on MANE transcripts, we excluded non-canonical transcripts, whose relevance may vary across tissues or disease contexts (74). Exploring the effects of stop-loss variants in non-canonical transcripts therefore represents an important area for future research.

The rapid advancements in protein language models and structure prediction tools, such as AlphaFold or ESM1b, offer promising opportunities for future model development (24, 75). Protein language models, which utilize vast datasets of protein sequences, effectively capture intricate patterns in amino acid composition and sequence features. Specifically, the functional impacts of in-frame indels have been well-predicted using ESM1b. Integrating these models into stop-loss variant prediction frameworks could greatly enhance our understanding of how extended proteins affect folding, stability, and function. Future iterations of TAILVAR could leverage these advanced tools to comprehensively evaluate the structural and functional consequences of stop-loss variants.

In conclusion, this study identifies key determinants of the pathogenic potential of stop-loss variants and proposes a systematic interpretive framework for their classification. Our findings show that transcript and protein features—particularly the properties of C-terminal extensions—are critical considerations in variant interpretation. The proposed framework and insights presented here have the potential to improve the interpretation of stop-loss variants and to guide future developments in the field. Consequently, TAILVAR can serve as a powerful tool for predicting the molecular effects of nonstop extensions, thereby enhancing the accuracy of genetic diagnoses and facilitating the discovery of novel disease-associated genes.

## Data availability

All code and data used to produce the TAILVAR model, figures, and results presented in this study are available in the GitHub repository at https://github.com/dr-yoon/TAILVAR. TAILVAR scores for all possible single-nucleotide substitutions and indels are pre-computed and can be downloaded from Zenodo (https://zenodo.org/records/16561299).

## Supplementary data

Supplementary Figures S1–S7. Supplementary Tables S1–S8.

## Acknowledgments

This research was supported by the Global Physician-Scientist Development Program – Postdoctoral Research Support for Early-Career Physician-Scientists grant (RS-2025-02214700, to JGY) from the Korea Health Industry Development Institute (KHIDI), funded by the Ministry of Health and Welfare, Republic of Korea.

## Conflict of of interest statement

The authors declare no conflicts of interest.

## Notes

### Competing Interest Statement

The authors have declared no competing interest.

