## Supplementary Figure for "A predictive framework for stop-loss variants with C-terminal extensions"

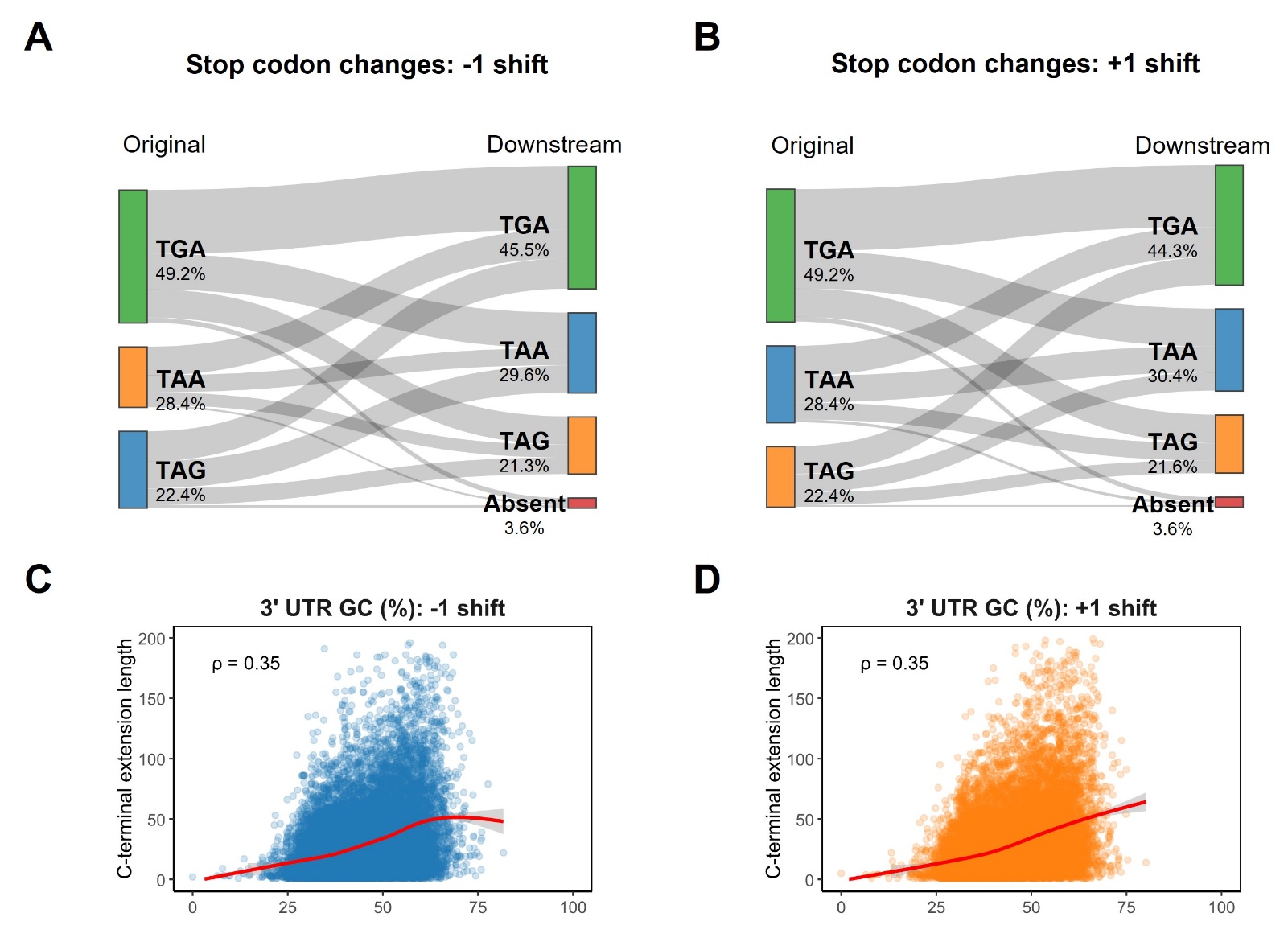

**Figure S1.** Stop codon usage is preserved following frameshift events.

(**A, B**) Sankey diagram illustrating changes in stop codon usages due to frameshifts (-1 shift and +1 shift) at the canonical stop codon site. Original stop codons (left) and their downstream equivalents after a -1 shift (**A**) or +1 shift (**B**) are indicated, with the width of the bands representing the proportion of transcripts undergoing each transition. Despite the frameshift, TGA remains the most frequently used stop codon. The frequency of TAA increases slightly, and overall stop codon usage patterns remain largely conserved. (**C, D**) Scatter plots showing the relationship between 3′ UTR GC content and the length of C-terminal extensions after a -1 shift (**C**) or +1 shift (**D**). Red lines indicate the regression fits, with Spearman’s correlation coefficient (ρ) shown in each panel. Points represent individual transcripts.

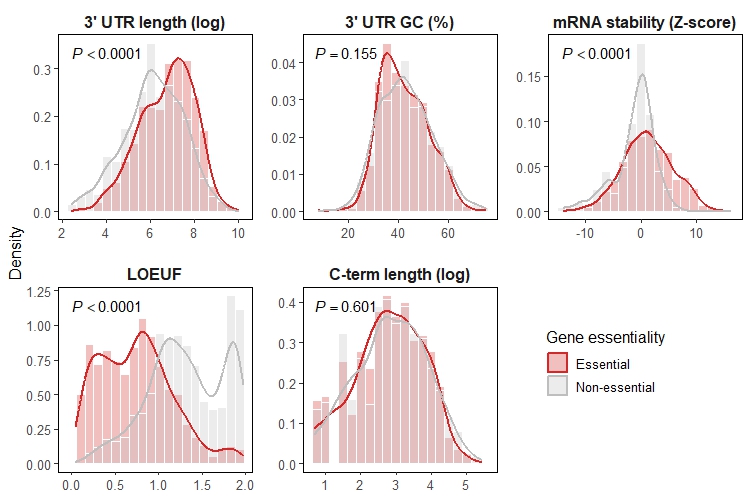

**Figure S2.** Transcript features differ by gene essentiality.

Density plots comparing transcript-level properties between essential (red) and non-essential (blue) genes. Features examined include 3′ UTR length (log-transformed), 3′ UTR GC content, mRNA stability (Z-score), LOEUF scores, and C-terminal extension length (log-transformed). Histograms represent the distribution of each feature, overlaid with smoothed density curves. *P* values are derived from the Mann–Whitney U test. Essential genes exhibit significantly longer 3′ UTRs and greater mRNA stability, along with markedly lower LOEUF scores, consistent with stronger selective constraint.

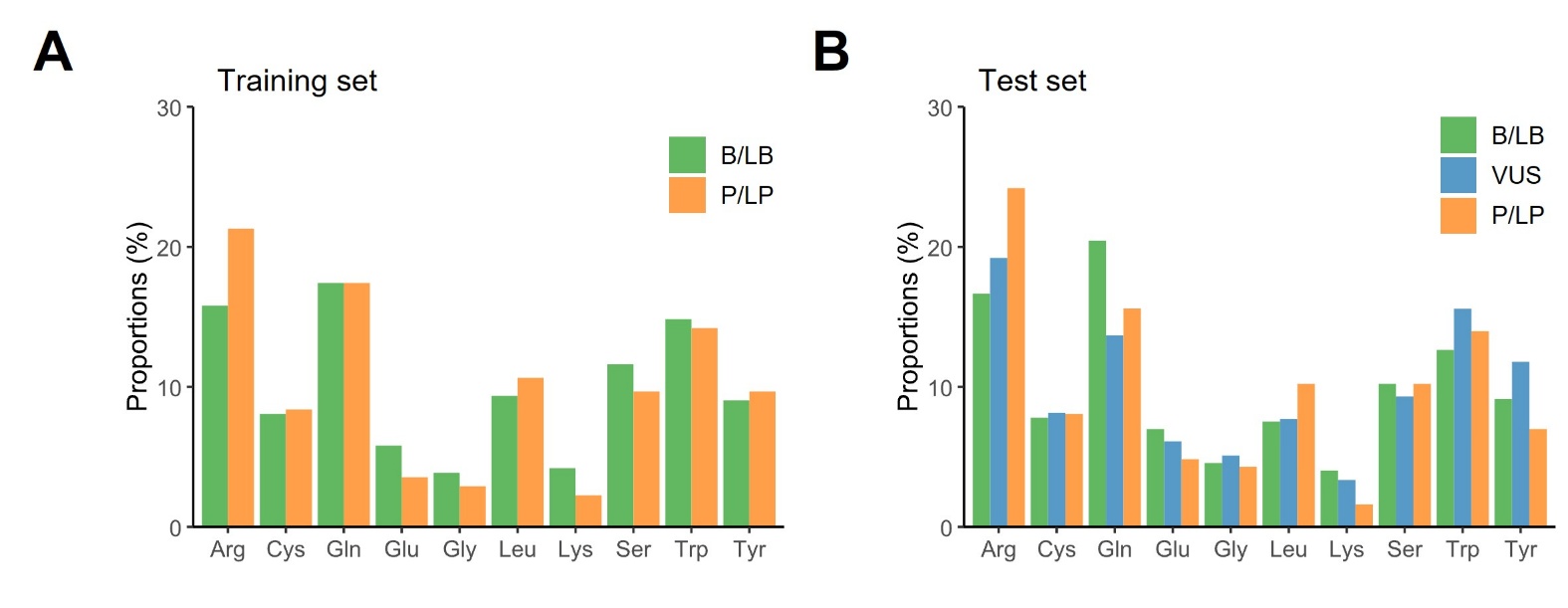

**Figure S3.** Distribution of amino acid substitutions resulting from stop‐loss variants.

Bar plots illustrate the proportional distribution of amino acid substitutions generated by single-nucleotide stop-loss variants, stratified by clinical pathogenicity classification. The analysis encompasses ten possible amino acid replacements, with bar height indicating the percentage of variants within each pathogenicity group. Variant categories are color-coded: benign/likely benign (B/LB; green), variant of uncertain significance (VUS; blue), and pathogenic/likely pathogenic (P/LP; orange). (**A**) Frequencies are shown for the training set; (**B**) frequencies for the independent test set. No specific amino acid substitution demonstrated a significant association with variant pathogenicity in either cohort.

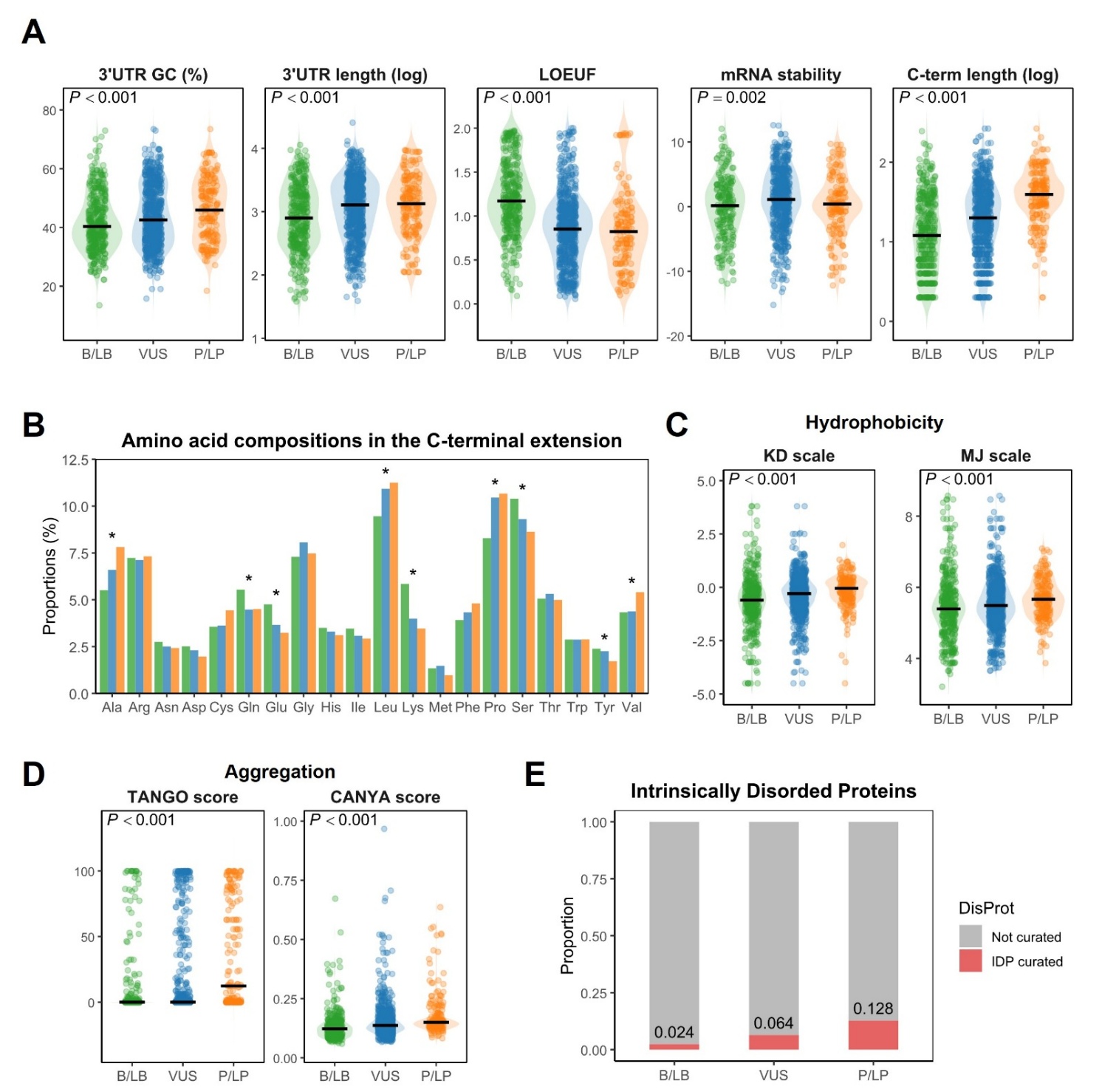

**Figure S4.** Transcript‐ and protein‐level characteristics of stop‐loss variants in the test set, stratified by clinical classification.

(**A**) Violin plots comparing transcript‐level features between B/LB (green), VUS (blue), and P/LP (orange) groups in the test set. From left to right: 3′ UTR GC content (percent), 3′ UTR length (log-transformed), LOEUF (LOF observed/expected upper bound), mRNA stability (Z‐score), and C‐terminal extension length (log-transformed). (**B**) Amino acid composition of aberrant C‐terminal extensions for B/LB (green), VUS (blue), and P/LP (orange) variants in the test set. Residues enriched in P/LP extensions (Ala, Cys, Leu, Pro) vs. B/LB extensions (Glu, Lys, Gln, Ser) are indicated by asterisks (* *P* < 0.05). (**C**) Hydrophobicity comparison of C‐terminal extensions using the Kyte–Doolittle (KD) and Miyazawa–Jernigan (MJ) scales. Violin plots show that P/LP‐derived peptides have significantly higher average hydrophobicity than B/LB peptides. (**D**) Proportion of intrinsically disordered proteins (IDPs) among genes harboring stop‐loss variants, based on DisProt annotation. P/LP variants (right) show a higher IDP fraction (13.1%) compared to B/LB variants (2.6%).

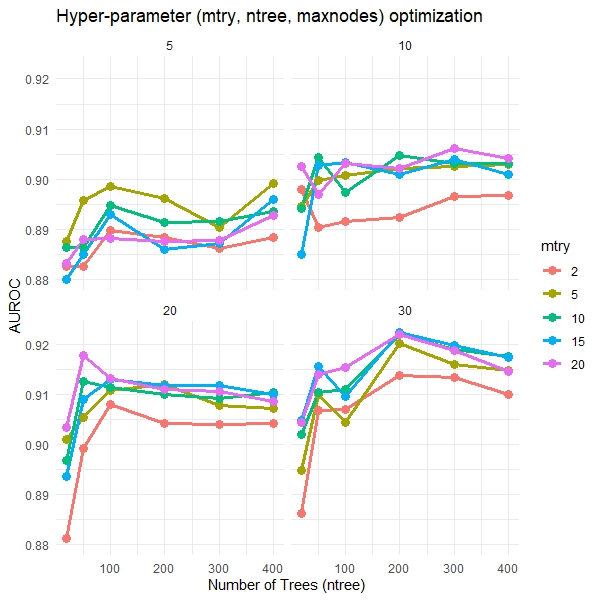

**Figure S5.** Hyperparameters optimization for TAILVAR model development.

To identify the optimal hyperparameters for the TAILVAR model, we conducted a comprehensive grid search over three critical parameters: *mtry* (number of variables randomly sampled as candidates at each split), *ntree* (number of trees in the forest), and *maxnodes* (maximum number of terminal nodes). Specifically, *mtry* was tested at values of 2, 5, 10, 15, and 20, *ntree* at 20, 40, 60, 80, and 100, and *maxnodes* at 5, 10, 20, and 30. The optimal combination, *mtry* = 20, *ntree* = 200, and *maxnodes* = 30, was selected based on maximizing the area under the receiver operating characteristic (AUROC) curve, a measure of classification accuracy.

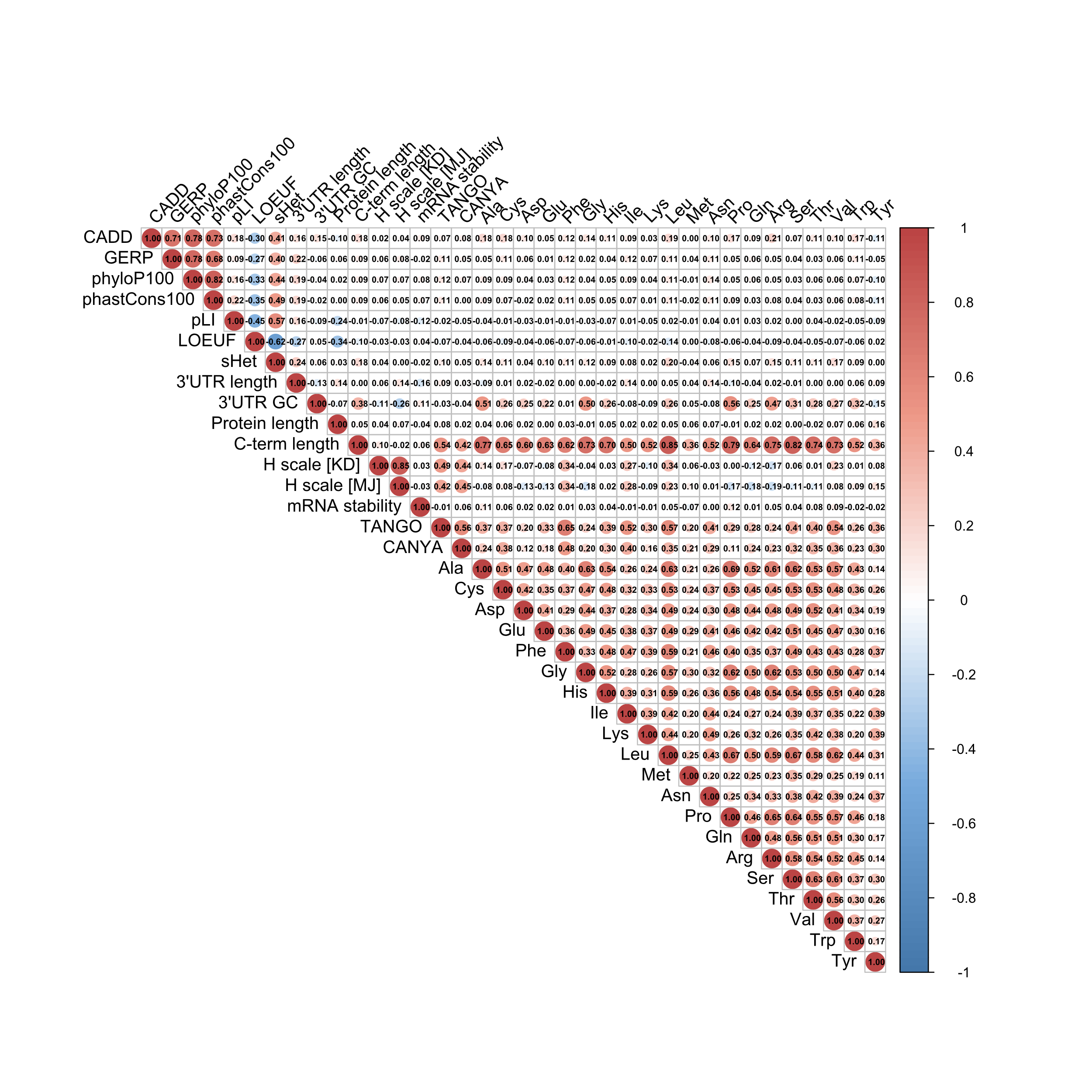

**Figure S6.** Correlation plot of features incorporated into the TAILVAR model.

The computational tools and conservation scores exhibit strong inter-correlations, whereas transcript and protein features show lower correlations with these tools. This distinction highlights the unique and complementary value of transcript and protein features in predicting the pathogenicity of stop-loss variants within the TAILVAR model.

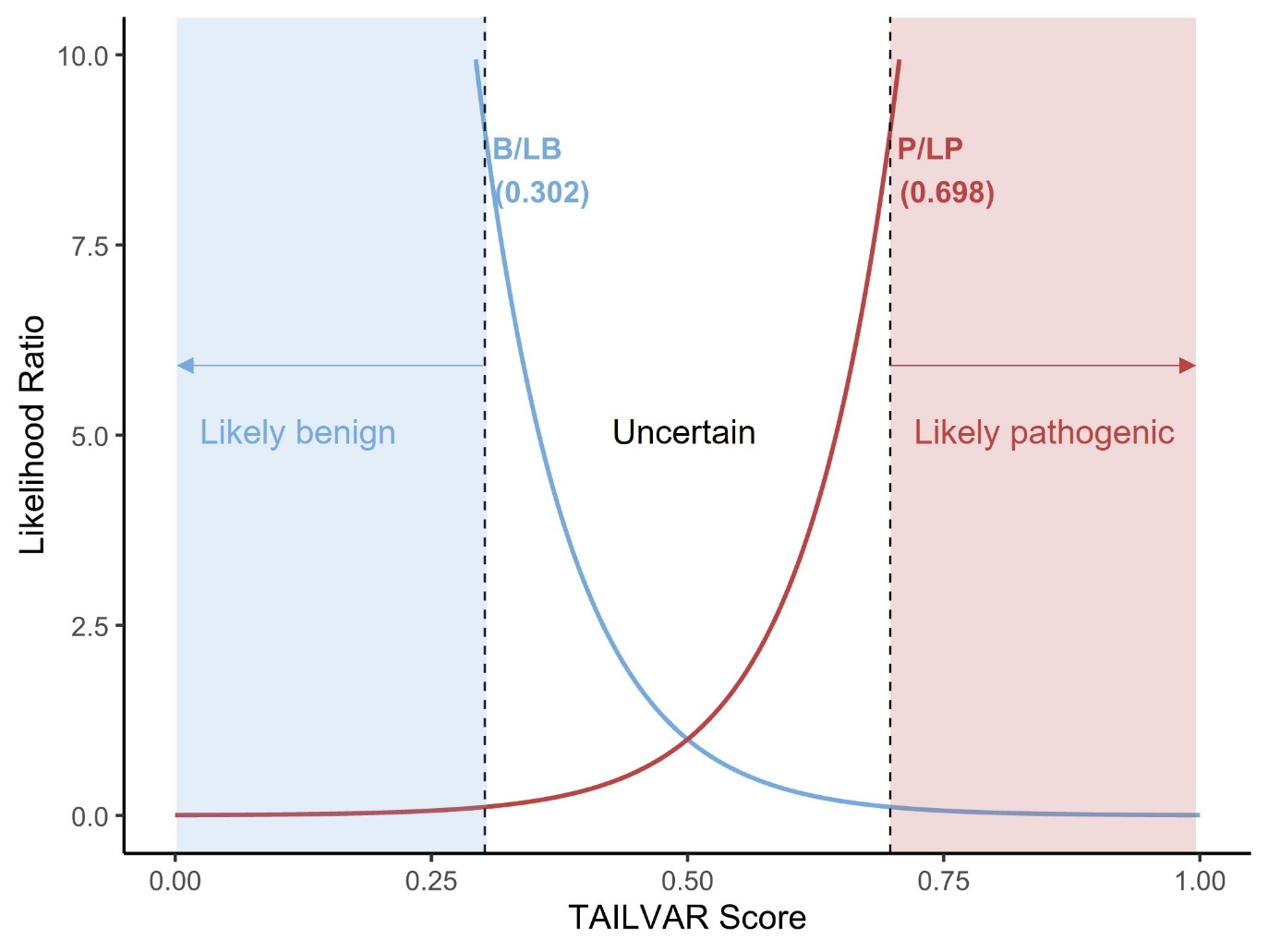

**Figure S7.** Determination of the TAILVAR score threshold.

A two-component Gaussian mixture model was fitted to the full spectrum of simulated single-nucleotide variants (*n* = 140,334) to estimate the distribution of TAILVAR scores for benign/likely benign (B/LB, blue) and pathogenic/likely pathogenic (P/LP, red) variants. Likelihood ratio (LR) curves were generated for both classes, and classification thresholds were defined as the TAILVAR scores corresponding to LR values of 9:1 for P/LP (right, red) and 1:9 for B/LB (left, blue), corresponding to a 90% posterior probability for each class. Arrows indicate the interpretation boundaries: variants with TAILVAR scores below the B/LB threshold are considered likely benign, those above the P/LP threshold are considered likely pathogenic, and those in between are classified as uncertain. Vertical dashed lines indicate the final thresholds for variant interpretation
